# Human neuron subtype programming through combinatorial patterning with scRNA-seq readouts

**DOI:** 10.1101/2023.12.12.571318

**Authors:** Hsiu-Chuan Lin, Jasper Janssens, Ann-Sophie Kroell, Philipp Hornauer, Malgorzata Santel, Ryoko Okamoto, Kyriaki Karava, Marthe Priouret, Maria Pascual Garcia, Manuel Schroeter, J. Gray Camp, Barbara Treutlein

## Abstract

Human neurons programmed through transcription factor (TF) overexpression model neuronal differentiation and neurological diseases. However, programming specific neuron types remains challenging. Here, we modulate developmental signaling pathways combined with TF overexpression to explore the spectrum of neuron subtypes generated from pluripotent stem cells. We screened 480 morphogen signaling modulations coupled with NGN2 or ASCL1/DLX2 induction using a multiplexed single-cell transcriptomic readout. Analysis of 700,000 cells identified diverse excitatory and inhibitory neurons patterned along the anterior-posterior and dorsal-ventral axes of neural tube development. We inferred signaling and TF interaction networks guiding differentiation of forebrain, midbrain, hindbrain, spinal cord, peripheral sympathetic and sensory neurons. Our approach provides a strategy for cell subtype programming and to investigate how cooperative signaling drives neuronal fate.

## Introduction

The advent of cell fate engineering, including the direct programming of iPSCs into diverse cell types, has revolutionized research in developmental biology, disease modeling, and therapy development (1). Pioneering transcription factors (TFs) such as neurogenin 2 (NEUROG2 or NGN2) and achaete-scute homolog 1 (ASCL1) play critical roles in initiating and regulating neurogenesis within vertebrate nervous systems, and their pioneering role is transferable in vitro to induce rapid neuron formation from pluripotency (2; 3; 4). NGN2-induced neuronal cells (iNs) were shown to exhibit excitatory glutamatergic properties within two weeks of in vitro differentiation (4). However, these neurons are heterogenous, with a mixture of central and peripheral nervous system-derived neuron identities (5). This diversity might be linked to NGN2’s naturally varied in vivo expression pattern across regions such as the forebrain, midbrain, hindbrain, spinal cord, and peripheral nervous system (6; 7; 8). It remains unclear how to guide the proneural function of TFs into each associated lineage.

In human brain development, neuronal diversity is stimulated by diverging morphogen gradients (9). This has inspired researchers to combine cell type-specific TF overexpression with the addition of extrinsic patterning factors to achieve regional specification (reviewed in (10)). For instance, combining NGN2 overexpression with dual SMAD and WNT inhibition was shown to generate forebrain neurons, while the addition of a SMAD inhibitor and a WNT agonist generated anterior hindbrain neurons (11; 12). However, there is a lack of systematic exploration of the neuronal diversity that can be achieved by combining TF overexpression and morphogen exposure and of systematic profiling of cellular heterogeneity within these induced cell populations.

Single-cell RNA sequencing (scRNA-seq) has emerged as a powerful tool for dissecting cellular heterogeneity and enhancing the resolution of cell identity (13). By examining the transcriptomes of individual cells during dynamic developmental processes, scRNA-seq enables the reconstruction of differentiation trajectories and the identification of underlying gene regulatory networks (GRNs) (14; 15; 16). This technique has been instrumental in creating comprehensive catalogs of human cells, which facilitates the characterization and validation of programmed cell fates and improves the selection of disease models (17). Moreover, the recent advances in scRNA-seq allows for sample multiplexing and the profiling of an increased number of cells (18; 19), providing exciting opportunities for high-throughput perturbation screening with readouts at single-cell level. In this study, we set out to systematically explore the neuronal cell type diversity that can be achieved using combinatorial morphogen patterning coupled with NGN2 or ASCL1/DLX2 induction. We used highly-multiplexed scRNA-seq readout to profile nearly 700,000 cells in 480 distinct morphogen combinations and found that iNs can be patterned into diverse neuron subtypes with identities aligned to the anterior-posterior and dorsal-ventral axes of neural tube development. Additionally, we developed a computational pipeline to analyze cooperative signaling and transcriptional regulation driving cell fate transitions, indicating the power of using in vitro systems to dissect morphogen response pathways. Taken together, our systematic screening approach provides valuable insights into neuron subtype specification and the underlying morphogen-dependent regulators, and it can serve as a general strategy for cell subtype programming.

## Results

### Combinatorial patterning expands the diversity of NGN2-iN subtypes

To systematically program neuron subtypes, we implemented a combinatorial morphogen screening strategy on NGN2-iNs from iPSCs. Our aim was to investigate whether the identity, concentration, and combination of morphogen signaling modulators could influence neuron subtype specification. Our selected compounds play pivotal roles in modulating signaling pathways crucial for patterning and regionalization in early nervous system development. These include four anterior-posterior (AP) morphogen signaling modulators: XAV-939 (XAV, a WNT pathway antagonist), CHIR-99021 (CHIR, a WNT pathway agonist), retinoic acid (RA), and FGF8 and three dorsal-ventral (DV) morphogen signaling modulators: BMP4, SHH, and purmorphamine (SHH pathway agonist). Employing different concentrations and combinations of these morphogen signaling modulators, we created a total of 24 AP patterning conditions and 8 DV patterning conditions, resulting in 192 unique AP x DV morphogen signaling modulator combinations for screening (Fig. S1A). For each of the combinations, we first induced NGN2 expression in iPSCs using doxycycline, followed by sequential exposure to corresponding AP morphogen signaling modulators and DV morphogen signaling modulators (Fig. 1A, Fig. S1A). This temporal exposure mimicked the natural developmental patterning timeline (20). After continuous exposure to morphogen combinations, NGN2-iNs from each condition were individually dissociated at day 10 (Fig. 1A), followed by nuclei isolation, fixation, permeabilization and processing for highly multiplexed single-nucleus RNA sequencing (snRNA-seq) using a split-pool combinatorial barcoding platform (ParseBiosciences).

**Figure 1.**
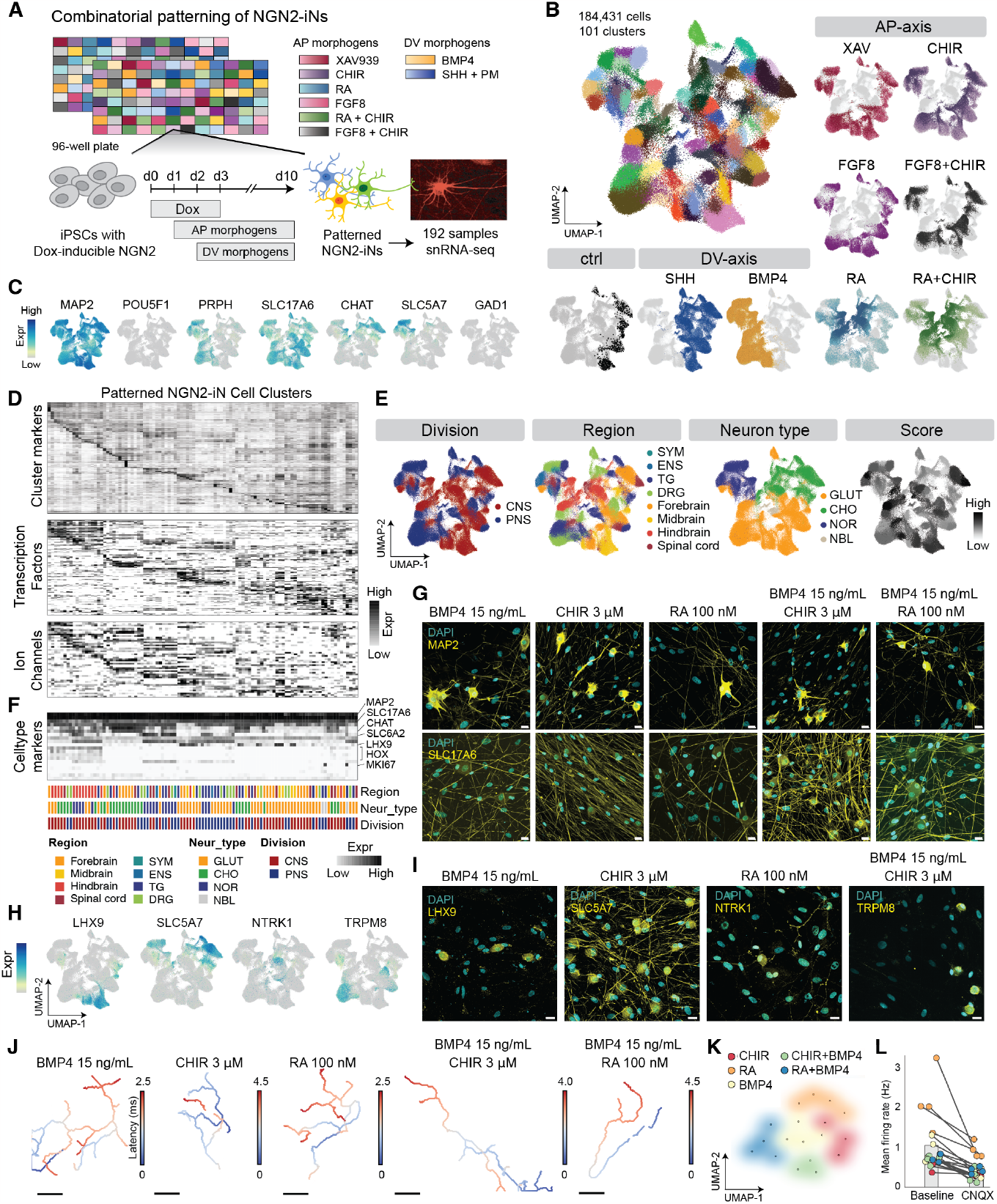
Combinatorial patterning screen to program NGN2-iN neuron subtypes. (A) Experimental scheme and timeline of the NGN2-iN combinatorial patterning screen with single-cell transcriptomics readout. NGN2-iNs exposed to each of the 192 morphogen combinations were individually analyzed with snRNA-seq using split-pool combinatorial barcoding (Parse Biosciences). (B) UMAP embedding of 184,431 cells in the dataset, colored by cluster identity and the source of AP- or DV-morphogens. (C) Feature plots of representative marker genes. (D) Heatmap of cluster markers, transcription factors and ion channel expression in each patterned NGN2-iN cell cluster. (E) UMAP embeddings colored based on the annotations transferred from primary neuron reference atlases, including division, region, neuron type and corresponding mapping score. CNS, central nervous system; PNS, peripheral nervous system; SYM, sympathetic nervous system; ENS, enteric nervous system; TG, trigeminal ganglia; DRG, dorsal root ganglia; GLUT: glutamatergic neuron; CHO, cholinergic neuron; NOR, noradrenergic neuron; NBL, neuroblast-like cells. (F) Heatmap of representative cell type markers in each patterned NGN2-iN cluster with transferred annotations as side bars. (G) Immunofluorescent staining of week 6 patterned NGN2-iNs with MAP2 (top row) and SLC17A6 (bottom row) (yellow). DAPI is shown in cyan. Scale bar: 20 µm. (H-I) Feature plots (H) and immunofluorescent staining (I) of neuron subtype-specific markers, including LHX9, SLC5A7, NTRK1 and TRPM8. Scale bar: 20 µm. (J) Example spike-triggered electrical footprints of NGN2-iNs recorded on high-density microelectrode arrays (HD-MEAs) for five patterning conditions. Colors indicate the latencies of action potentials propagating along the various neurites (from blue to red). Scale bar: 200 µm. (K) Clustering of iN patternings could be clustered based on the electrophysiological features obtained from HD-MEA recordings (each dot represents one neuronal network). (L) CNQX application significantly reduced the spontaneous electrical activity of iNs across all patterning conditions; coloring as in (K).

Following data filtering to exclude low-quality cells and potential multiplets, we successfully profiled 184,431 cells across 192 conditions, with an average of 1000 cells per condition (Fig. S1B, Supplementary Table 1). Our analysis revealed 101 molecularly distinct clusters of patterned NGN2-iNs (Fig. 1B, see Methods). We utilized Uniform Manifold Approximation and Projection (UMAP) embedding to visually represent the dataset. Notably, the morphogen signaling modulator treatments significantly increased the diversity of NGN2-iNs (Fig. 1B). A pronounced separation and mirrored cluster distribution was observed between BMP4-treated and SHH-treated NGN2-iNs, recapitulating their distinct roles in DV patterning and neural fate specification. Canonical marker gene expression analysis indicated that patterned NGN2-iNs primarily exhibited a neuronal identity (MAP2+), transitioning away from pluripotent stem cell states (POU5F1-) and ceasing cell cycle activity (MKI67-) (Fig. 1C). The majority of patterned NGN2-iNs are glutamatergic neurons (SLC17A6+), while there are also cholinergic neuron (CHAT+) and noradrenergic neuron (SLC6A2+) subtypes present (Fig. 1C). Nearly half of the patterned NGN2-iNs expressed peripherin (PRPH+), a typical marker for the peripheral nervous system (PNS), and the expression of PRPH correlated strongly with BMP4 treatment (Fig. 1B-C). We provided a detailed list of cluster marker genes, including TFs and ion channels (ICs) that are among the differentially expressed genes (DEGs) for each molecularly distinct cluster (Fig. 1D).

We annotated the patterned NGN2-iNs by transferring labels from a diverse set of human and mouse primary single cell transcriptomic atlases of fetal and adult nervous systems encompassing all major brain regions, retina, spinal cord, enteric nervous system (ENS), dorsal root ganglia (DRG) and trigeminal ganglia (TG) (21; 22; 23; 24; 25; 26; 27; 28; 29) (Supplementary Table 2, see Methods). To enhance robustness in label transfer, we employed high-resolution clustering by aggregating similar cells within the same cluster (Fig. S1C). These high-resolution clusters faithfully represented the overall cluster distribution and significantly expanded the number of features available for analysis (Fig. S1C to E). Patterned NGN2-iNs are mapped to a spectrum of distinct neuron types spanning the anteriorposterior axis of central nervous system development, including forebrain, midbrain, hindbrain, and spinal cord regions (Fig. 1E). Beyond the central nervous system (CNS), certain clusters of patterned NGN2-iNs exhibited regional identities associated with the PNS, including the sympathetic nervous system, enteric nervous system, trigeminal ganglia, and dorsal root ganglia (Fig. 1E). Our mapping is supported by relevant cell type markers, including the presence of the LHX9 hgene in midbrain clusters and HOX genes in hindbrain clusters (Fig. 1F). We validated the expression of MAP2 and SLC17A6 as well as neuron subtype-specific marker genes LHX9, SLC5A7, NTRK1 and TRPM8 in selected samples using immunostaining (Fig. 1G to I, see Methods). To probe the functionality of patterned NGN2-iNs, we performed electrophysiological measurements by means of high-density microelectrode arrays (HD-MEAs). Patterned NGN2-iNs developed spontaneous electrical activity early on (Fig. S2, Fig. S3, Movie S1), and demonstrated robust axonal outgrowth (Fig. 1J, Movie S1). Moreover, we found that different patterning conditions manifest in distinct electrophysiological activity (Fig. 1K), and that this activity could be significantly reduced by the AMPA/kainate receptor antagonist 6-Cyano-7-nitroquinoxaline-2,3-dione (CNQX) (Fig. 1L; Fig. S3). These findings underscore the efficacy of combinatorial patterning in inducing diverse neuron subtypes from NGN2-iNs.

### Morphogen combinations activate distinct gene regulatory networks that shape neuronal diversity in patterned NGN2-iNs

To understand how morphogen signaling modulators and their combinations drive diverse NGN2-iN sub-types, we analyzed the diversity of clusters that are generated by each condition (Fig. 2A). We found that all conditions give rise to multiple clusters, suggesting that combinatorial patterning is insufficient to generate homogeneous neuron subtypes (Fig. 2A to B, Fig. S4A). Yet, we were able to identify a preference of morphogen signaling modulator combinations to drive NGN2-iNs towards certain nervous system regions or neuron types (Fig. S4B to C). Notably, we found that for XAV, FGF8, BMP4 and SHH, the identity of the signaling modulator is dominant in cluster specification, while the concentration contributes only to a minor extent (Fig. 2A). Conversely, for RA the concentration has a dramatic effect on cluster identity, and with increasing RA concentration cells gradually shift to new cell clusters (Fig. 2A to B). The addition of BMP4 to the RA treatment yielded a pronounced acquisition of new cell states, indicating the presence of a synergistic effect between BMP and RA (Fig. 2B). Similar cooperative effects are also observed for combinatorial treatment with BMP4 and CHIR, while no synergy is observed between SHH and RA (Fig. 2B, Fig. S5A). Next, we quantified the number of clusters per condition, which showed a strong concentration-dependent effect of RA on cell fate restriction, independent of SHH or BMP4 (Fig. 2B to D). Conversely, CHIR induces increased cellular heterogeneity at medium concentrations, but a drop in cell cluster diversity at the highest concentration (Fig. 2B to D). BMP4 in general tends to produce fewer cell clusters compared to both the control (ctrl) and SHH conditions, yet the combinatorial effect of BMP4 and a high concentration of XAV leads to an intriguing increase in the number of clusters (Fig. 2C).

**Figure 2.**
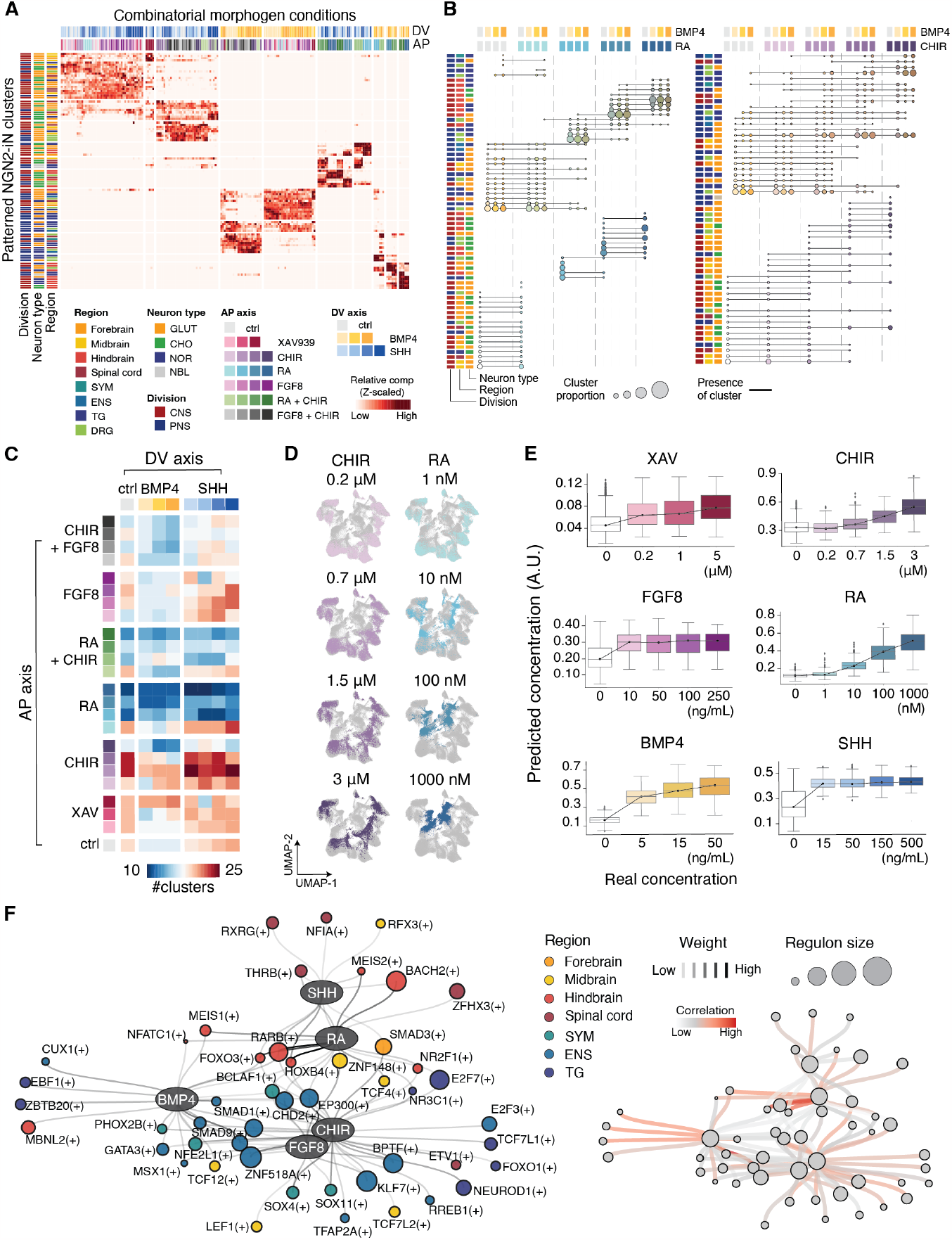
Dissecting the regulatory mechanisms of combinatorial morphogen-induced neuronal diversity. (A) Heatmap visualization of the contribution of each combinatorial morphogen condition to patterned NGN2-iN clusters. Cell count in each morphogen condition is normalized for comparison. Morphogen conditions and annotations of cell clusters are color labeled as side bars. (B) Dot plot illustrating cell composition and identity shift in response to morphogen combinations. Each column represents a distinct morphogen combination, while each row corresponds to a cluster with color-coded annotations along the side. The dots label the presence of a cluster, with their size denoting the proportion within each morphogen combination. Connections via lines indicate the continuity of a cluster across different morphogen combinations. (C) Number of cell clusters generated in each morphogen combination. (D) UMAP embedding with cells colored by CHIR and RA gradients. (E) Actual versus predicted morphogen modulator concentrations from gene sets identified from random forest classifier of morphogen responses. (F) Gene regulatory network (GRN) centered on morphogen signaling modulators. Left: each oval node represents a morphogen modulator, linked to circular nodes indicating associated regulons (TF and its target genes). Edge colors indicate the strength of the association between morphogen signaling modulators and regulons. The regulons are color-coded based on regional identity where the regulons are maximaly active. The size of the nodes corresponds to the size of the regulons, indicating the number of target genes regulated by the TF. Right: The same GRN where edge color represents the correlation of regulon activity with morphogen modulator concentration.

We looked into how morphogen signaling modulators affect the identity of NGN2-iN subtypes. Notably, RA treatment results in more NGN2-iNs with hindbrain cholinergic neuronal identity, in line with the known function of RA signaling in posterior patterning and cholinergic neural induction (Fig. 2A-B, Fig. S4) (30; 31). WNT inhibition resulted in a subtle enhancement of forebrain fate, while its activation reduced the proportion of forebrain and midbrain cells. Activation of WNT also induced cells with distinct TG and DRG regional characteristics, accompanied by increased proportions of cholinergic and noradrenergic neuronal identity, consistent with existing research findings (Fig. S4) (32; 33; 34; 35; 36; 37). SHH treatment exhibited a dosage-dependent effect on anterior pattern acquisition as well as cholinergic neurogenesis, aligning with previous research (Fig. S4A, S4C, Fig. S5A) (38; 39). Conversely, BMP4 drove an increase in noradrenergic neuronal identity and a general regional specification within the PNS as reported previously (Fig. 2A to B, Fig. S4A, S4C) (40; 41; 42; 43; 44). We did not observe any significant effect by FGF8 treatment.

We used two approaches to elucidate the underlying molecular events orchestrated by the morphogen signaling modulators. We first trained a random forest-based regression model to predict morphogen concentrations based on gene expression data, with model performance as a proxy to quantify the difference between morphogen concentrations. We found that SHH and FGF8 responses reached saturation even at the lowest concentrations (Fig. 2E). In contrast, BMP4 and XAV exhibited minor dosage effects (Fig. 2E). RA and CHIR displayed the most robust dosage-dependent effects, with response saturation yet to be achieved even at the highest concentrations (Fig. 2E). Next, we investigated the genes used by the model and found synergistic molecular features in the interplay between RA and BMP4 as well as CHIR and BMP4, which mirrored the effect observed in the acquisition of new cell clusters (Fig. 2B, Fig. S5B to C). Conversely, SHH treatment together with RA did not induce a strong combinatorial molecular shift (Fig. S5A, S5D).

To explore the regulatory mechanisms governing patterned NGN2-iNs fate specification in more detail, we inferred GRNs with SCENIC (15). We identified regulons, transcription factors and their respective target genes, associated with specific morphogen signaling modulators. We found that each morphogen modulator was linked to distinct sets of regulons that exhibited enriched activity in specific nervous system regions, with some regulons being controlled by multiple morphogen signaling modulators (Fig. 2F, Fig. S6A). RA-associated regulons displayed a pronounced enrichment in hindbrain fates, including well-known drivers of posterior patterning such as HOXB4, RARB, and MEIS (Fig. 2F, Fig. S6A to B) (45; 46). FGF8-associated regulons had the propensity to promote midbrain fates (Fig. 2F, Fig. S6A). BMP4 and CHIR-associated regulons exhibited strong links to the peripheral nervous system (Fig. 2F). Notably, known effectors of BMP4, such as PHOX2B, SMAD1, SMAD9, and MSX1, were also identified within these regulons (Fig. 2F) (47; 48). Many of the aforementioned regulons exhibited positive activity correlation with the concentrations of their respective morphogen (Fig. 2F, Fig. S6B). Additionally, we calculated specifically activated regulons within each molecular cluster (Fig. S6C). PHOX2B and TLX2 regulons are highly associated with noradrenergic neuron fates, while THRB and RXRG were linked with cholinergic neuron fates, in line with previous research (Fig. S6C) (49; 50; 51; 52). These findings suggested that combinatorial morphogen exposure activates distinct transcription factor regulons that shape the identity and diversity of patterned NGN2-iN cells.

### NGN2 induction in diverse pre-patterned neural progenitor cell input states leads to neuronal identities distinct from post-patterned NGN2-iNs

While our study has shown that combinatorial patterning following NGN2 induction (post-patterning) from iPSCs can induce diverse neuron subtypes (Fig. 1), patterning and regionalization typically precedes neuronal differentiation in natural neural development (53). We therefore assessed whether the timing of patterning influences the specification of programmed neural subtypes, and whether diverse input states affect or expand the repertoire of NGN2-iN types. We set up a pre-patterning screen where we first patterned iPSCs into distinct regionalized neural progenitor cells followed by NGN2 induction (Fig. 3A). We used three different basal media to induce a neural lineage during pre-patterning, including general neural induction media (NIM), dual SMAD inhibition favoring anterior patterning (N2B27-2Si) and SMAD inhibition coupled with WNT activation promoting posterior patterning (N2B27-SB-CHIR). Following one day of neural induction, diverse AP and DV patterning conditions were provided to the cells, resulting in a total of 96 pre-patterning conditions that served as input states for NGN2 induction (Fig. 3A, Fig. S7). We phenotyped these regionalized neural progenitor states using single-cell multiomic analysis, including both transcriptome and open chromatin modalities, combining all 96 samples into one pool (Fig. 3A to B, Fig. S8). In addition, we collected bulk RNA-seq data for each individual pre-patterned sample, which allowed us to demultiplex the sc-multiome data into the 96 conditions (Fig. 3A, Fig. S8).

**Figure 3.**
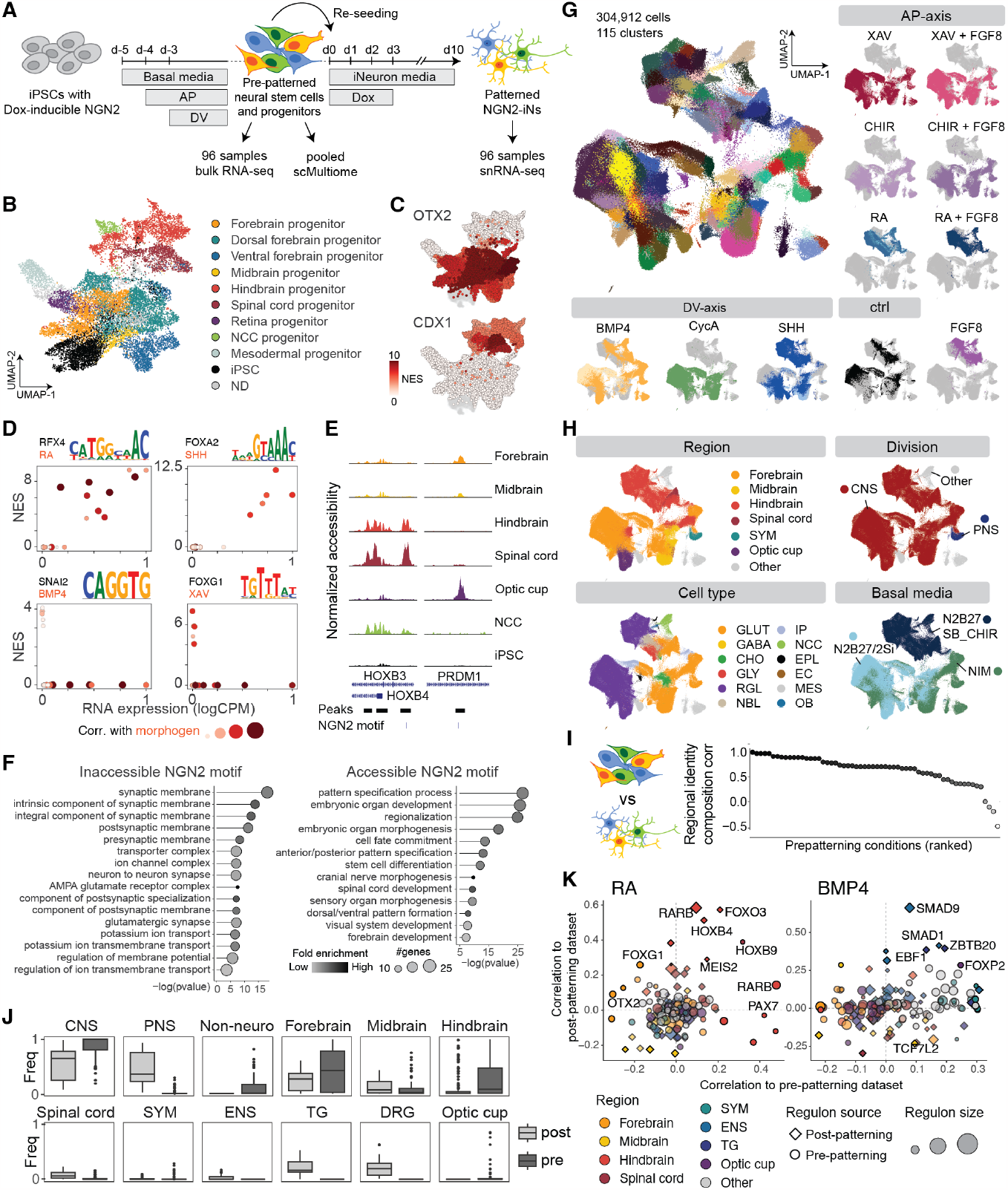
Inducing NGN2 from diverse pre-patterned input states expands programmed neuronal diversity. (A) Experimental scheme and timeline for the pre-patterning screen coupled with NGN2 induction. iPSCs were pre-patterned into diverse neural stem cells and neural progenitors followed by NGN2 induction. Pre-patterned neural stem cells and neural progenitors from 96 conditions were harvested individually for bulk RNA-seq as well as pooled for single-cell multiome (10x Genomics). The resulting pre-pattered NGN2-iNs were analyzed individually with snRNA-seq. (B) UMAP embedding of pre-patterned neural stem cells and neural progenitors colored by annotation. NCC, neural crest cells; iPSC, induced-pluripotent stem cells; ND, not determined. (C) UMAP embedding colored by motif enrichment in accessible regions of that cluster (normalized enrichment score, NES). (D) Scatter plot showing the relationship between motif enrichment and TF expression. Positive correlations are linked to activators, negative correlations indicate chromatin closing factors (repressors). (E) Example of ATAC peaks showing differential accessibility of NGN2-motifs. (F) Gene ontology (GO) analysis for genes linked with inaccessible or accessible NGN2-motifs. (G-H) UMAP embedding of 304,912 cells in the pre-patterned NGN2-iN dataset, colored by cluster identity, the source of AP- or DV-morphogens (G), the annotations transferred from primary neuron reference atlases and the basal media used (H). GABA, GABAergic neuron; GLY, glycinergic neuron; RGL, radial glia-like cell; IP, intermediate progenitor; EPL, ependymal-like cell; EC, endothelial cell; MES, mesenchymal cell; OB, osteoblast. (I) Neural identity composition correlation between pre-patterned neural stem cells and the resulting pre-patterned NGN2-iNs. Each dot represents a morphogen combination condition. (J) Comparison of annotated divisions and regional identity emerged from pre- and post-patterned NGN2-iNs shown as composition (frequency: freq.). (K) Comparison of regulons inferred from pre- and post-patterned NGN2-iNs and their correlation with RA and BMP4. Regulons inferred separately from pre- and post-patterned datasets represented by different shapes were scored for activity correlation to indicated morphogen gradients in both datasets. Regulons are colored with the regional identity where maximal activity is registered, while the size of the regulons was represented as the size of the dots.

Computational analysis of the sc-multiome data of 96 pre-patterned samples revealed a spectrum of diverse neural progenitor cells (NPCs) exhibiting distinct regional identities (Fig. 3B). The chromatin landscape showed drastic changes upon exposure to different morphogen concentrations and/or basal media, with thousands of regions altering in accessibility (Fig. S9). Motif enrichment on differential morphogen patterning conditions reveals major players controlling regional neural fate including OTX2 in forebrain progenitors (54) and CDX1 in hindbrain progenitors (55) (Fig. 3C). Combining gene expression data with motif enrichment also reveals functional roles for transcription factors, which we could link to morphogen signaling modulator concentrations. We found that RFX4 expression is high in clusters linked to RA, and additionally RFX4’s motif is highly enriched in regions accessible in cells expressing RFX4, suggesting an activating role (Fig. 3D). Similarly, the activator FOXA2 was linked to SHH, consistent with previous research (56) (Fig. 3D). Repressing factors, where expression of a factor leads to closing of regions with its motif, were also found, for example for BMP4 (SNAI2) and XAV (FOXG1) (Fig. 3D). Next, we assessed the accessibility of NGN2 motifs in the various regionalized NPC states. We filtered the NGN2 motifs based on chromatin accessibility to identify potential NGN2 target genes associated with these peaks, and analyzed the enrichment of biological processes (Fig. 3E to F). Inaccessible NGN2 motifs are involved in neural maturation, suggesting that, following induction, NGN2 as pioneer factor might open up these regions to induce neuronal identity (Fig. 3F). In contrast, accessible NGN2 motifs are associated with patterning and regionalization processes, exhibiting differential accessibility across distinct pre-patterned progenitors (Fig. 3E to F, Fig. S9).

Next, we wondered how these diverse regionalized NPC input states with distinct chromatin accessibility landscapes would respond to NGN2 induction. We therefore induced NGN2 expression in all conditions using doxycycline and phenotyped the resulting cells using highly multiplexed snRNA-seq, analogous to the experimental procedure applied in the post-patterning screen (Fig. 3A, Fig. S7). We successfully profiled 304,912 cells across 96 conditions and identified 115 molecularly distinct clusters (Fig. 3G). The majority of clusters exhibited a neuronal identity including neuron-like, but interestingly also less mature radial glia-like cells, and we observed a small number of non-neural off-target cells (Fig. 3G). Pre-patterned NGN2-iNs exhibit diverse regional and cellular identity distinct from control (Fig. 3G to H, Fig. S10). We compared the regional identity composition of regionalized NPC input states and corresponding NGN2-iNs and found that regional identity is largely preserved (Fig. 3I, Fig. S9). This implies that in most cases, NGN2 is not actively involved in regionalization but rather induces neuronal differentiation from the regional identity pre-established in progenitors.

We compared pre-patterned and post-patterned NGN2-iNs to assess their similarities and to identify new cell states acquired using the pre-patterning regime. We found that pre-patterned and post-patterned cells acquire different region identity composition and cell types (Fig. 1E, Fig. 3H, Fig. 3J). Cells in the pre-patterning condition displayed a higher proportion of neuronal cells of the CNS and an increase of off-target non-neuronal cells compared to the post-patterning condition (Fig. 3J). Enrichments in forebrain and hindbrain regional identity are observed, presumably due to the specific basal media preferences used during pre-patterning (Fig. 3J). Retinal progenitors are highly enriched in dual SMADi/WNTi/BMP4 conditions, whereas neural crest cells are enriched in SMADi/WNTa/BMP4 conditions (Fig. S10). Higher concentrations of BMP4, SHH, and XAV appeared to elevate the proportion of off-target non-neuronal cells (Fig. S10). These findings demonstrate that the pre-patterning regime with NGN2 induction in regionalized NPCs indeed provides inroads to new neuronal identities beyond those obtained during post-patterning.

We investigated the gene regulatory mechanisms driving these cell fate differences (Fig. S11). We inferred the GRNs of pre-patterned NGN2-iNs and compared the regulons to that of post-patterned NGN2-iNs (Fig. 3K, Fig. S11). Morphogen signaling modulators generally activate distinct regulon networks in both pre- and post-patterned NGN2-iNs, with some shared regulons like MEIS2, HOXB4, and HOXB9 associated with RA, as well as SMAD1, EBF1, and ZBTB20 associated with BMP4 (Fig. 3K, Fig. S11). Intriguingly, while the RARB regulons in pre- and post-patterning are both associated with RA and hindbrain regionality, their activities show low correlation with RA when assessed in the alternate dataset (e.g., pre-regulon score in post-dataset and vice versa) (Fig. 3K). This suggests that although the RARB transcription factor is induced by RA, it regulates different sets of target genes in pre- and post-patterned NGN2-iNs. Our findings underscore the distinct effects of NGN2 induction in the pre-patterning versus post-patterning regimes.

### Leveraging TF induction, combinatorial patterning and scRNA-seq readout as a general strategy for cell subtype programming

To assess the potential of combinatorial patterning as a general strategy for cell subtype programming, we extended our investigation to ASCL1-DLX2-induced GABAergic neuronal cells (ASCL1/DLX2-iNs). Adopting a similar experimental framework used for combinatorial patterning coupled with NGN2 induction, we designed 192 different combinations of AP and DV morphogen signaling modulators for pre- and post-patterned cells coupled with ASCL1/DLX2 induction. We performed snRNA-seq on all conditions and profiled 85,756 and 139,711 cells, respectively (Fig. 4A to B, Fig. S12 to Fig. S16). Computational analysis identified 78 and 107 clusters for pre- and post-patterned cells, respectively, and revealed a GABAergic neuronal phenotype for a large fraction of the cells indicated by expression of MAP2 and GAD1, and the absence of expression of SLC17A6. Similar to the findings with NGN2-iNs, we observed a higher proportion of off-target cells without MAP2/GAD1 expression in the pre-patterning compared to the post-patterning conditions (Fig. 4B). We combined data from all patterned NGN2- and ASCL1/DLX2-induced cells to establish an integrated induced neuronal cell atlas (Fig. 4C). Co-clustering all samples revealed minimal overlap between the datasets, indicating the efficacy of our strategies in generating diverse neuronal subtypes (Fig. 4C). To explore the differences in morphogen responsecurves between NGN2- and ASCL1/DLX2-iNs, we quantified the similarity between morphogen signaling modulator conditions and controls in relation to the exposed modulator concentration (Fig. 4D). Notably, ASCL1/DLX2-iNs are more sensitive to SHH concentrations, showing more pronounced changes in gene expression primarily at intermediate concentrations (Fig. 4D). This aligns with previous research showing a higher sensitivity of GABAergic neurons to ventralizing signals (57). Apart from SHH, we find that the other morphogens follow similar dynamics in post-patterned NGN2- and ASCL1/DLX2-iNs (Fig. 4D). Interestingly, while response to XAV reached a plateau in both post-patterned NGN2- and ASCL1/DLX2-iNs, we find that the highest concentration of XAV affects post-patterned NGN2-iNs to a greater extent (Fig. 4D). Moreover, we identified distinct interactions between basal media and morphogen dynamics in pre-patterned cells independent of the induced TF (Fig. S12). This observation emphasizes the distinct action of morphogens in various input states represented by different basal media. It also underscores the potential of distinct pioneering TFs in driving differential gene expression changes from a similar input state, which may arise from their differential interaction with established epigenetic states in prepatterned neural progenitors.

**Figure 4.**
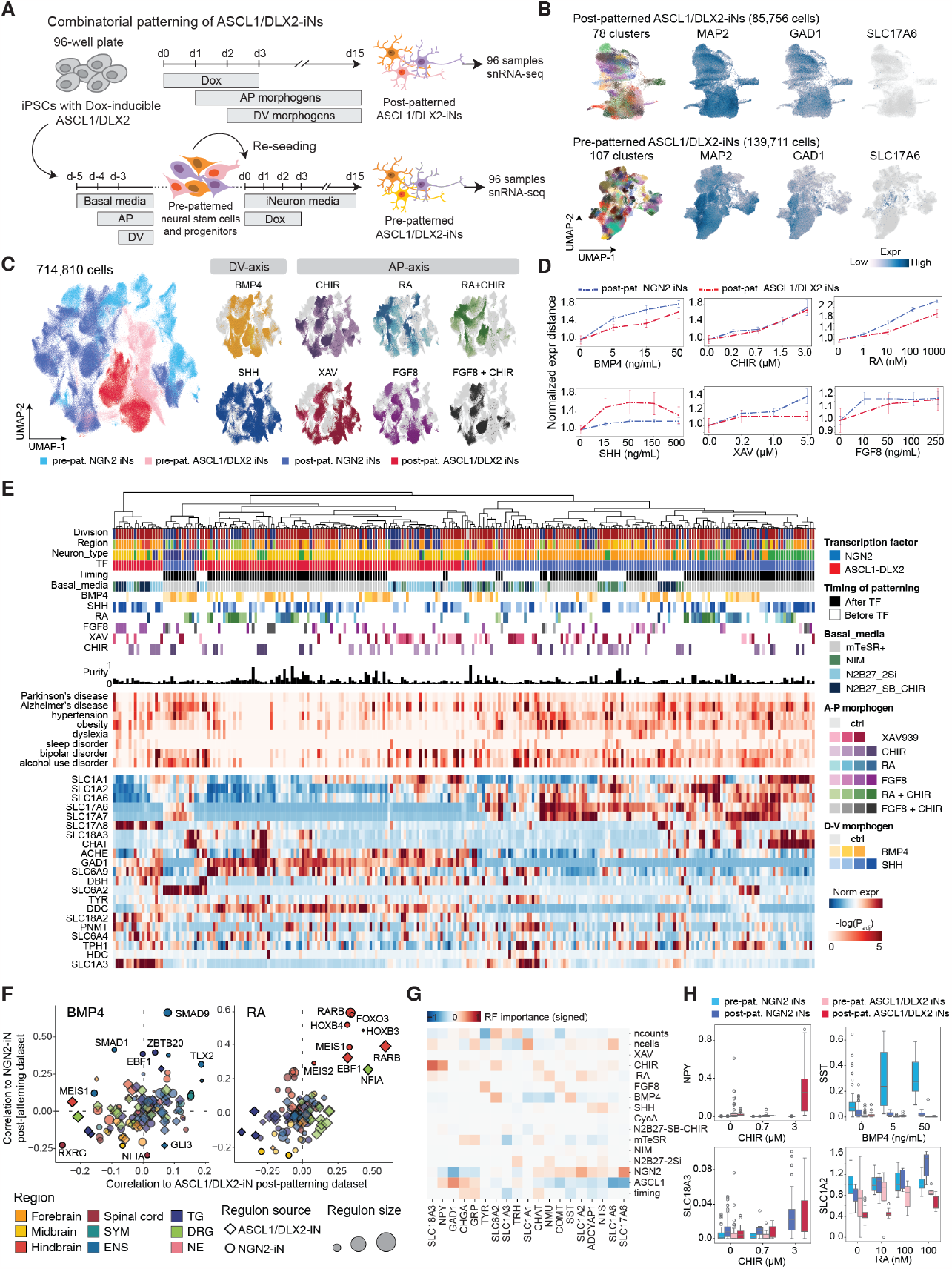
Combinatorial patterning generates diverse GABAergic ASCL1/DLX2 iNs. (A) Experimental scheme and timeline for the pre- and post-patterning screen coupled with ASCL1/DLX2 induction. (B) UMAP embedding of pre-and post-patterned cells after ASCL1/DLX2 induction. ASCL1/DLX2-iNs are mostly neuronal (MAP2+) and GABAergic (GAD1+), with no or low expression of excitatory markers (SLC17A6-). (C) UMAP embedding of integrated atlas of all neuronal cells (n=430,936 cells) showing sample distribution and concentration gradients. (D) Scatterplots showing mean distance to control cells (normalized to 1) in relation to morphogen concentrations. Error bars represent SEM. (E) Diversity of clusters generated in all datasets, shown by dendrogram based on transcriptomic distance (cosine, TF marker genes). Annotation heatmap shows concentrations for the condition which has the highest relative composition of the cluster. Purity shown in percentage as barplots. GO heatmap shows enrichment of disease genes in marker genes of clusters. Expression heatmap shows row-normalized expression of key neurotransmitter genes, highlighting the diversity. (F) Comparison of regulons inferred from post-patterned NGN2-iNs and ASCL1/DLX2-iNs and their correlation with BMP4 and RA. Regulons inferred separately from NGN2- and ASCL1/DLX2 datasets represented by different shapes were scored for activity correlation to indicated morphogen gradients in both datasets. Regulons are colored with the regional identity where maximal activity is registered, while the size of the regulons was represented as the size of the dots. (G) Heatmap showing importances of morphogens for a Random Forest regressor to predict expression of neurotransmitter and neuropeptide genes. Importances were multiplied with the sign of the Pearson correlation to indicate directionality. (H) Boxplots showing examples of neurotransmitter/neuropeptides correlated with morphogen concentrations. The box plots show the median (center line), upper and lower quartiles (box limits) and 1.5×interquartile range (whiskers). Outliers shown.

To assess the similarity between all identified cell clusters, we performed clustering based on TFs and ion channels (Fig. 4E). We associated each cluster with the condition that exhibited the highest proportion of cells within that specific cluster. Our analysis revealed an initial cluster segregation based on timing (pre- versus post-patterning) and TFs, followed by the influence of the basal media and combinatorial morphogen conditions (Fig. 4E). The purity of clusters that can be reached under optimal conditions ranges from 0.3% to 84%, with a mean of 15%. 71 clusters can be generated with over 25% purity (Fig. 4E). To understand the similarity of morphogen response in each combinatorial patterning scheme, we systematically compared the morphogen-centered regulons across datasets. Our findings revealed substantial similarity between the regulons of post-patterned NGN2-iNs and ASCL1/DLX2-iNs (Fig. 4F, Fig. S17). This suggests a shared impact of morphogens on TF-induced neurons post-TF induction, despite variations in proneuronal TFs.

iNs exhibit promising potential for studying diseases in vitro, serving both fundamental and pharmaceutical research purposes (10). To assess the potential of patterned iNs in disease modeling, we used signatures from Human Disease Ontology (58) to perform gene set enrichment on the marker genes of each cluster and identified several clusters with enriched expression of disease-related genes. Patterned iNs also display expression of a wide repertoire of different neurotransmitters (NTs) and neuropeptides (NPs). Using Random Forest modeling, we were able to link certain morphogen concentrations and conditions to NT/NP expression changes (Fig. 4G). We find that the highest concentration of CHIR is associated with GABAergic neurons that express neuropeptide Y (NPY), a neuropeptide that has been linked to obesity and alcohol abuse (Fig. 4H) (59). Furthermore, increased concentrations of BMP4 in pre-patterned NGN2-iNs is associated with higher expression of somatostatin (SST) (Fig. 4H). Post-patterned NGN2-iNs exposed to higher RA concentrations, on the other hand, show elevated expression of the glutamate transporter SLC1A2 that is known for its association with various neuropathologies (Fig. 4H) (60). These findings suggest the potential of patterned iNs in simulating pathological conditions.

## Discussion

Human neurons generated from iPSCs using defined TFs offer exciting opportunities in disease modeling and therapeutics screening using human cells. This approach is transformative because of the high induction efficiency and its rapid maturation compared to in vivo neurons, significantly expediting research progress. Despite the potential of iNs to generate brain regionspecific neuron subtypes by coupling with morphogen signaling (11; 12), a systematic exploration of morphogen combinations with iNs and the resulting achievable neuronal diversity has been lacking. In this study, we investigated how morphogen signaling modulator identity, gradients, combination, and treatment timing cooperate with different pioneering TFs to influence neuronal differentiation across 480 conditions. Using high-throughput scRNA-seq as a readout, we individually profiled each condition, successfully capturing a total of 700,000 cells and identified over 400 molecularly distinct clusters. Drawing upon extensive scRNA-seq primary neuron atlases (21; 22; 23; 24; 25; 26; 27; 28; 29), we mapped the patterned iNs, annotated their identities, and assessed their similarity to primary counterparts. We showed that patterned iNs cover diverse neuron subtypes that are mapped across a range of regionality along the neural axis with various neurotransmitter identities. This programmed neuronal diversity atlas can serve as a rich resource to facilitate further engineering of desired neuron subtypes.

iNs have certain limitations. We previously found that NGN2-iNs are molecularly heterogeneous (5), and it has been challenging to match these neuron populations to their primary counterparts with high confidence. While combinatorial patterning successfully expands the diversity of induced neuronal cells, resulting patterned iNs remain molecularly heterogeneous. While we have identified several new conditions that can achieve 80 % neuronal subtype homogeneity (Fig. 4E), most iN cultures remain heterogeneous. Additional morphogen signaling modulators or overexpression of different TFs, might help to improve specificity. We identified several regional and neurotransmitter subtype-specific regulons, and these TFs could serve as a starting point in further engineering of homogeneous sub-types with desired features. Currently, our annotation of patterned iNs are based on their relative similarity to primary neurons. Although we observe a preference towards specific regionality compared to unpatterned controls, their transcriptomes are still not identical to those of primary neurons. Considering absolute similarity, there remains a considerable distance between patterned iNs and in vivo neurons. This discrepancy could result from their TF-induced nature and the presence of a iN gene expression background. Eliminating this background represents a key obstacle in enhancing the similarity of iNs to primary neurons.

All together, our data represents an important foundation in programming iPSCs into specific neuronal cell types. We performed a large-scale combinatorial screen of morphogen signaling modulators, activating and inhibiting important developmental pathways. We have shown how these pathways interact with each other, and their molecular underpinnings in the form of GRNs to create neuronal cell types. We present strategies for the programming of distinct cell types with specific neurotransmitter, TF expression and disease relevances, and our methodology opens up the possibility for future work to enlarge the repertoire of iNs.

## ACKNOWLEDGEMENTS

We thank all members of the Treutlein and Camp labs for discussions and feedback. We thank Fátima Sanchís-Calleja for insightful discussion on morphogens, patterning, brain regionalization and for sharing reagents. We thank Jonas S. Fleck for insightful discussion on GRN inference, regulons and high resolution clustering. We thank Zhisong He and Tomás Gomes for general bioinformatics discussion and support. We thank Christian Beisel for advice on Illumina sequencing. Illumina sequencing was performed at the Genomics Facility Basel at D-BSSE, ETH Zurich. We thank the Microscope facility at D-BSSE, ETH for training and technical support. We thank Oliver Brüstle, Michael Peitz and Universitatsklinikum Bonn for sharing the ASCL1-DLX2-inducible iPSCs. H.-C.L. is supported by a Human Frontier Science Program fellowship (LT000399/2020-L) and an EMBO long term fellowship (ALTF 1190-2019). J.J. is supported by an EMBO long term fellowship (ALTF 32-2023). M.P., M.P.G and M.Sc. were supported by an Innosuisse grant (101.366 IP-LS). This work was supported by the European Research Council (Braintime-874606 (B.T.)), the Swiss National Science Foundation (Project Grants 310030_192604 (B.T.), 310030_84795 (B.T. and J.G.C.)), and the National Center of Competence in Research Molecular Systems Engineering (B.T.).

## AUTHOR CONTRIBUTIONS

H.-C.L generated and cultured induced neurons used in this study. H.-C.L. generated all single-cell transcriptome datasets with support from M.Sa, and R.O.. J.J. and H.-C.L. performed bioinformatics analysis and visualization of single-cell transcriptome and single-cell multiome data. J.J., H.-C.L., B.T. and J.G.C. interpreted the results. A.-S.K. performed the pre-test and assisted in NGN2-iN pre-patterning screen data collection. H.-C.L and A.-S.K. generated the single-cell multiome data and bulk RNA-seq data of pre-patterned neural progenitors with support from M.Sa.. M.Sa. performed immunostaining and analyzed the data with support from H.-C.L.. K.K. generated the single-cell multiome data of iPSCs and established iGABA culture with support from H.-C.L.. M.P., M.Sc. and M.P.G. generated the electrophysiological data. P.H. analyzed and visualized the electrophysiological data. M.Sc. supervised the electrophysiological experiment. H.-C.L., B.T. and J.G.C. designed the study and H.-C.L., J.J., A.-S.K., B.T. and J.G.C. wrote the manuscript.

## Material and Methods

### iPSC culture

All cells were cultured under standard conditions of 37°C and 5% CO2 unless specified otherwise. NGN2-inducible iPSCs, originating from the 409B2 background (RIKEN BRC Cell Bank), were generated following previously established protocols (5). ASCL1-DLX2-inducible iPSCs, sourced from the iLB-C-133bm-s4 background (Universi-tatsklinikum Bonn, https://hpscreg.eu/cell-line/UKBi013-A) (61), were acquired from Prof. Dr. Oliver Brüstle’s research group at Universitatsklinikum Bonn under an arranged Material Transfer Agreement (MTA). NGN2-inducible iPSCs were cultured in mTeSR Plus Medium (STEMCELL Technologies), whereas ASCL1-DLX2-inducible iPSCs were sustained in StemMACS iPS-Brew medium (Miltenyi Biotec). Both iPSC lines were maintained on matrigel-coated plates under feeder-free conditions using respective media. To sub-culture iPSCs, cells were treated with TrypLE (Gibco) and then incubated overnight in media supplemented with 5 µM of the Rho-associated protein kinase (ROCK) inhibitor Y-27632 (STEMCELL Technologies).

### NGN2-iNs induction, culture and combinatorial patterning

For post-patterning, NGN2-inducible iPSCs were dissociated into a single-cell suspension using Accutase (STEM-CELL Technologies) and plated on 50 µg/mL poly-L-ornithine (PLO, Sigma-Aldrich) and 10 µg/mL Laminin from mouse Engelbreth-Holm-Swarm sarcoma (LMN, Sigma-Aldrich)-treated plates in mTeSR Plus media with 5 µM ROCK inhibitor. NGN2 expression was induced using 4 µg/mL doxycycline (Sigma-Aldrich). The day after NGN2 induction (day 1), the medium was changed to DMEM/F-12 (Gibco) supplemented with 4 µg/ml doxycycline, 1:100 N-2 supplement (Gibco), 1:100 MEM nonessential amino acid solution (NEAA, Gibco), 10 ng/µL human BDNF (Pepro-Tech), 10 ng/µL human NT-3 (PeproTech), 5 µM ROCK inhibitor, 0.2 µg/mL laminin, and corresponding AP patterning morphogens (Supplementary Table 1). On day 2, corresponding DV patterning morphogens were applied on top of AP patterning morphogens without changing of media (Supplementary Table 1). From day 3 onwards, combinations of AP and DV patterning morphogens were supplemented in Neurobasal medium (Gibco) with 1:50 B-27 supplement (Gibco), 1:100 GlutaMAX supplement (Gibco), 10 ng/µL human BDNF, 10 ng/µL human NT3, and 1 µM Cytosine-D-arabinofuranoside (Ara-C, Sigma-Aldrich) to eliminate proliferating cells. Doxycycline was removed after day 3. At days 5, 7, and 9, NGN2-iNs were maintained in the same media as on day 3 but without Ara-C. Patterned NGN2-iNs were collected for snRNA-seq analysis on day 10.

For pre-patterning, NGN2-inducible iPSCs were seeded into three distinct types of basal media supplemented with 5 µM ROCK inhibitor, including:

1. NIM: DMEM/F12, 1:100 N-2 supplement, 1:100 Glutamax, 1:100 MEM-NEAA, 1 µg/mL Heparin, and 1:200 Penicillin-Streptomycin (Gibco).
2. N2B27-2Si: 50% DMEM/F12, 50% Neurobasal, 1:100 N-2 supplement, 1:100 B27-VtA, 1:100 ITS-G, 1:100 Glutamax, 1:200 Penicillin-Streptomycin, 10 µM SB-431542 (Miltenyi Biotec), and 100 nM LDN-193189 (STEM-CELL Technologies).
3. N2B27-SB-CHIR: 50% DMEM/F12, 50% Neurobasal, 1:100 N-2 supplement, 1:100 B27-VtA, 1:100 ITS-G, 1:100 Glutamax, 1:200 Penicillin-Streptomycin, 10 µM SB-431542, and 3 µM CHIR99021 (Tocris Bioscience).

One day after seeding (day 1), AP patterning morphogens were introduced into the respective basal media (Supplementary Table 1) during a media change. By day 2, DV patterning morphogens were directly added into the existing media containing corresponding AP morphogens. At day 3, a media change was performed alongside the addition of both AP and DV patterning morphogen signaling modulators. Following five days of pre-patterning, prepatterned neural stem cells and progenitors were dissociated using Accutase. For each pre-patterning condition, NGN2-iNs were induced and cultured using the aforementioned procedures but without Ara-C. In parallel, cells from each pre-patterning condition were harvested for bulk RNA-seq as well as pooled for 10x multiome analysis.

### iGABA induction, culture and combinatorial patterning

iGABA were induced and cultured using a previously established protocol (62). For post-patterning, ASCL1-DLX2-inducible iPSCs were dissociated using Accutase and plated on matrigel-coated plates containing iPS-Brew supplemented with 5 µM ROCK inhibitor. The following day, media were changed to remove ROCK inhibitor. Media were changed again the next day (day 0) into neural induction media (DMEM/F12 + 1:100 N-2 supplement) containing 2 µg/mL doxycycline to induce ASCL1-DLX2 expression. Doxycycline remains in the media until day 8. At day 2, cells were dissociated into single-cell suspension using accutase and seeded on PLO/LMN-coated plates with neuronal media (NBB27, composed of Neurobasal, 1:50 B27 supplement, 1:100 Glutamax, 10 ng/mL human BDNF, 1:500 LMN) containing 2 µg/mL doxycycline and 5 µM ROCK inhibitor. The media were changed at day 3 to remove the ROCK inhibitor and to add 10 µM DAPT (Tocris Bioscience), which remained until day 7. On day 4, the media were changed using NBB27 media containing doxycycline and DAPT, supplemented with AP patterning morphogens (Supplementary Table 1). On day 5, DV patterning morphogens were added directly on top of corresponding AP patterning morphogens without media change (Supplementary Table 1). At day 6, media were changed into neuronal media containing AP and DV morphogen combinations as well as 1 µM Ara-C and 1:500 LMN. At day 8, 10, 12 and 14, media change were performed using the same media composition containing AP and DV morphogen combinations but without Ara-C, DAPT and doxycycline. Patterned iGABAs were collected for snRNA-seq analysis on day 15. Pre-patterning of iGABA was conducted following procedures similar to those outlined for NGN2-iNs, utilizing the iGABA neuronal induction and culture conditions.

### Single-nuclei isolation, fixation, snRNA-seq library preparation, sequencing and data pre-processing

To obtain single-cell suspensions, induced neurons cultured in PLO/LMN-coated 12-well plates were washed with PBS and incubated with 300 µL accutase at 37°C for 5 mins. Dissociation reactions were stopped with 700 µL PBS containing 0.04% BSA, and cells were gently triturated into single-cell suspensions. Cells were washed twice with 500 µL PBS containing 0.04% BSA before nuclei isolation. To isolate the nuclei from single-cell suspension, cells were triturated ten times in 100 µL lysis buffer (10 mM Tris-Cl pH7.4, 3 mM NaCl, 10 mM MgCl2, 1% BSA, 0.1% Tween-20, 1 mM DTT, 0.2U/µL RNase inhibitor (Takara), 0.1% NP-40, 0.01% Digitonin (Invitrogen)) and incubated 2 mins. on ice. The isolated nuclei underwent two subsequent washes in 500 µL wash buffer (10 mM Tris-Cl pH7.4, 3 mM NaCl, 10 mM MgCl2, 1% BSA, 0.1% Tween-20, 1 mM DTT, 0.2U/µL RNase inhibitor). Isolated nuclei were fixed and cryopreserved for snRNA-seq according to the instructions provided by the manufacturer (Parse Biosciences). snRNA-seq library preparation, sequencing, and initial data pre-processing were strictly adhered to the guidelines provided by the manufacturer (Parse Biosciences). Notably, pair-end single index sequencing parameters were adjusted to 101/6/0/86 (as opposed to 74/6/0/86) to effectively utilize the capabilities of the NovaSeq 200 cycle reagent kit (Illumina). To demultiplex each sample and to generate the matrix of transcript counts in each cell, sequencing reads from fastq files were aligned against the human genome and transcriptome reference (hg38, sourced from Parse Biosciences) by the split-pipe (v1.0.4, Parse Biosciences) analysis pipeline using default parameters.

### scRNA-seq data processing

All scRNA-seq samples were processed individually. First, count matrices obtained by split-pipe were loaded into Scanpy. Next, cells were filtered on quality using the following thresholds: minimum number of genes = 750 and minimum number of UMI counts = 1000. Furthermore, cells that were more than 5 standard deviations in log1p(total_counts), log1p(number of genes) and percentage of counts in top 20 genes were also removed. Next, we removed cells that showed too high percentage of mitochondrial reads (5% for NGN2-iNs (pre and post-patterned), 6% for ASCL1/DLX2 post-patterned iNs, 8% for ASCL1/DLX2 pre-patterned iNs). Finally, we used Scrublet to remove predicted doublets. This led to a final cell count of 184,431 post-patterned and 304,912 pre-patterned NGN2-iNs; 85,756 post-patterned and 139,711 pre-patterned ASCL1/DLX2 iNs.

We filtered out genes that were expressed in fewer than 3 cells per sample and the data was log normalized using 10,000 as a scaling factor. Highly variable genes were selected using default parameters in Scanpy (min_mean=0.0125, max_mean=3, min_disp=0.5), and the data was then subset and scaled (maximum value of 10). Effects of total counts were regressed out, since the first clustering showed large effects. PCA was performed and significant PCs were selected based on the variance ratio, leading to 80 PCs for NGN2-iNs pre-patterned, ASCL1/DLX2 pre and post-patterned. NGN2-iNs post-patterning were sequenced in two runs, which were first integrated using Harmony. 60 PCs were selected for downstream analysis.

Clustering was performed using the Leiden algorithm, using resolution 10. This leads to a high-resolution clustering of the data. We established a script to merge similar clusters. First, average expression profiles were calculated per cluster. Next, differential gene expression was performed between every cluster and its most similar cluster (based on spearman correlation of the averaged profiles). Clusters were kept separate if at least 5 differential genes were detected (pval adj<0.05, lfc>2) of which at least 3 had to be functionally related (transcription factors, ion channels). After merging, the process was repeated until all clusters satisfied the constraints. This strategy led to 115 (NGN2-iNs, pre-patterned), 101 (NGN2-iNs, post-patterned), 107 (ASCL1/DLX2-iNs, pre-patterned) and 78 (ASCL1/DLX2-iNs, post-patterned) clusters.

### Annotation of scSeq data

To annotate the scSeq data, we performed high-resolution clustering within each cluster and aggregated the data from cells in each high-resolution cluster (highres cluster) into meta-cells. Subsequently, we performed canonical correlation analysis (CCA)-based label transfer (implemented in Seurat) (63), transferring labels from individual primary neuron reference atlases (Supplementary Table 2) onto the highres clusters. This label transfer was based on three gene sets: the 5,000 highly variable genes from our dataset, the highly variable genes from the reference atlases (normally encompassing around 30% of the detected genes), and cluster marker genes from our dataset. We determined the consensus results among the three gene sets and corresponding maximal matching scores within each highres cluster, and subsequently summarized these results to each cluster. The final cluster annotation was assigned based on consensus agreement among primary references, the percentage of highres clusters in each transferred category in each cluster, matching scores, and the expression of canonical marker genes. Each cluster was annotated by multiple layers of identities, including division (CNS, PNS, and non-neuronal), region, neuron type, and associated TF/ion channels defining that specific cluster (Supplementary Table 3).

### Morphogen RF classifier

To create random forest regressor models to predict morphogen concentrations based on gene expression data, we first subset our data to only contain conditions with at least 250 cells. We then subsampled all conditions to 250 cells. Next, we lognormalized morphogen concentrations and performed min-max normalization. The expression matrix used raw UMIs. The data was split into a train and test data set using train_test_split (test_size=0.11, shuffled). A RandomForestRegressor object was initialized with (n_estimators=100, max_features=‘sqrt’, max_depth=10) which was used in MultiOutputRegressor. This model was then used to predict concentrations on the test data. Every model was trained 5 times with different random_states and all functions used are part of the scikit-learn library.

For every morphogen we selected the top 150 genes with an importance score>0.001 in each of the 5 runs and merged the set. We then used hierarchical clustering of the averaged expression profiles of every morphogen concentration to create modules.

### Gene regulatory network analysis

Gene regulatory networks were inferred using SCENIC (15). We loaded the latest motif databases (motifs-v10, both 10kbp_up_10kbp and 500bp_up_100bp_down) and all human transcription factors from cistarget. We ran SCENIC 20 times on each dataset. Each iteration, we subsampled cells to acquire 100 cells per cluster (with replacement, using different random states each run). SCENIC was run using the pySCENIC implementation of GRNBoost2 and default parameters. The 20 runs were then used to create consensus regulons, only keeping TF-target gene interactions detected in at least 5 runs.

We used AUCell to score all cells based on the consensus regulons. We then used GRNBoost2 to link morphogen concentrations and basal media to regulon activity. We also hardcoded interactions in the matrix (var1*var2*var3) to better find interactions between morphogens and basal media. If a variable was linked to a target gene, but the link of an interaction variable had a higher weight than the individual variable, only the strongest link was kept and annotated as such. Finally, we performed filtering on the modules: w>250.

### scMultiome library preparation, sequencing and data pre-processing

Single-cell suspensions obtained from each NGN2-iN pre-patterning condition were divided into two distinct pools (Supplementary Table 1). Subsequently, each pool, along with the control NGN2-inducible iPSCs, underwent nuclei isolation individually following provider’s instruction (10x Genomics user guide CG000365). These isolated nuclei were then separately processed for 10x Multiome GEM generation, library preparation, and sequencing according to the provider’s instructions (10x Genomics user guide CG000338). To generate the matrices of transcript and fragment counts per cell, sequencing reads from the fastq file were aligned against the human genome and transcriptome reference (hg38, sourced from 10x Genomics) utilizing the Cell Ranger ARC (v2.0.0, 10x Genomics) analysis pipeline with default parameters.

### scMultiome data processing

The three RNA libraries were merged together and processed in Scanpy. Cells with fewer than 400 genes or more than 10,000 genes were filtered out. In addition, cells with more than 20% mitochondrial reads and over 50,000 UMIs were removed. Scrublet was used to predict and remove doublets.

Clustering was performed with the same parameters as in the other samples. Harmony was used for integration and 60PCs were used for subsequent analyses. Leiden clustering 4 was used as the high resolution clustering due to a lower number of cells compared to the other analyses. Clusters were merged if fewer than 5 differential genes were detected (pval adj<1e-10, lfc>2) with its most similar cluster or if fewer than 3 transcription factors or ion channels were differentially expressed.

ATAC libraries were processed in SCENIC+ starting from the fragment files. The RNA clustering was used to create pseudobulks, from which peaks were called using MACS2 (shift=73,ext_size=146,q_value=0.05,keep_dup=‘all’). Next, the get_consensus_peaks function was used to create a set of consensus peaks (peak width = 500bp). The compute_qc_stats function was used to calculate quality metrics (duplicate rate, TSS enrichment, FRIP score). We used the following filtering on cells: log(unique fragemens)>3.3, FRIP>0.45 and TSS enrichment>5.

To find variable regions, we first imputed accessibility (scale factor = 1,000,000) which we then normalized to a scale factor of 10,000. The function find_highly_variable_features was then used with default parameters. Differentially accessible regions were calculated using find_diff_features on imputed accessibility and variable regions, using the RNA clustering. Further, we calculated differential accessibility between every cluster and the control clusters. PycisTarget was used to calculate Motif enrichment on all differential regions using the motif 10 database.

### Bulk RNA-seq library preparation, sequencing and data pre-processing

To extract RNA, cells from each NGN2-iN pre-patterning condition were individually collected and lysed in Trizol (Invitrogen). Subsequently, RNA extraction was performed in the multi-well format using the MagMAX™-96 for microarrays total RNA isolation kit (Invitrogen). Quality assessment of the isolated RNAs from each condition was conducted using the RNA 6000 Pico Kit and the 2100 Bioanalyzer Systems (Agilent), while quantification was carried out using the Qubit system (Invitrogen). cDNA reverse transcription and amplification were carried out according to the smart-seq2 protocol (64). The preparation of the sequencing library was based on the Nextera XT DNA Library Prep protocol (Illumina), and subsequently sequenced using the Illumina NextSeq High Output paired-end 2×75 parameter. Sequencing reads from the fastq file were aligned against the hg38 reference genome (obtained from 10x Genomics) using the STAR RNA-seq aligner with default parameters.

### Demultiplexing of scMultiome from bulk RNA-seq

The MuSiC package was used to demultiplex bulk samples to single-cell clusters (65). In short, markers were calculated for every cluster using Seurat FindAllMarker (logfc>1, min.pct = 0.1). Then, music_prop was used to estimate cluster proportions (Leiden 0.25) in each bulk sample, using the marker genes. A low-resolution clustering was used since we found the algorithm unable to differentiate between smaller clusters. Instead we focused on linked larger clusters. We then calculated correlations between cluster proportions and morphogen/basal media conditions.

### NGN2-motif analysis

To obtain the NGN2-motif-containing fragments from the scMultiome ATAC data, we utilized the Pando gene regulatory network inference package (16) that has implemented the human base GRN (NetworkRegions@motifs@data) containing NGN2-binding motifs (MA0669.1, MA1642.1). ATAC fragments were extracted based on the presence of NGN2-binding motifs and subsequently filtered by accessibility. Fragments were considered accessible if detected in over 10% of the cells; otherwise, they were categorized as inaccessible. The accessible and inaccessible fragments were associated with genes based on their genomic proximity using rGREAT (66). Subsequently, the associated genes were analyzed for gene ontology enrichment using clusterProfiler (67).

### Distance response-curves

To obtain a quantitative effect of morphogen concentrations on gene expression changes, we calculated for every cell in each main experiment (pre/post, NGN2/ASCL1+DLX2) the euclidean distance in PCA space (50 components) to all control cells using the cdist function in scipy. We then averaged the distance over the 10 closest control cells and used this as a metric of gene expression change (the distance is affected both by cell type changes and up/down regulation of genes). Next, we calculated the average distance of control cells to other control cells in the same manner and used the mean control-control distance to normalize all other distances.

### Neurotransmitter/Neuropeptide linking

We used all neuronal clusters (based on MAP2, STMN2 and DCX and TUBB3 expression) and calculated averaged expression profiles for every condition (unique combination of AP-axis, DV-axis, timing, NGN2 or ASCL1/DLX2 and basal media). Basal media and timing were one-hot encoded and all variables were standardized. The expression matrices were log-normalized (scale factor 10,000). We trained a Random Forest Regressor model (n_estimators=100, max_depth=3) to predict expression of neurotransmitter and neuropeptide genes (expressed in > 100 cells, minimum expression level of 0.1) based on conditional variables. The model was scored using R-squared values, and only models with R2>0.1 were used. A second model was trained using OLS regression on the same input. The OLS feature importances were used to give signs to the RF feature importances to indicate directionality.

### Electrophysiology

Neuronal cultures were recorded with CMOS-based high-density microelectrode array (HD-MEA) recording systems, produced by MaxWell Biosystems (Zurich, Switzerland). In the present study, a 24-multiwell plate was used, comprising 24 HD-MEAs, each consisting of 26,400 low-noise electrodes with a center-to-center electrode pitch of 17.5 µm, arranged in a 220 × 120 electrode array structure. The used HD-MEA can record simultaneously from a total of up to 1020 user-selected readout-channels at 10 kHz sampling rate; the sensing area is 3.85 × 2.10 mm2. After 9 days in culture, patterned NGN2-iNs were dissociated into a single-cell suspension using Accutase (STEMCELL Technologies) and co-cultured with rat cortical astrocytes (Genlantis) on PLO/LMN-coated HD-MEAs (20,000 neurons + 40,000 astrocytes). Weekly recordings of the network activity started 12 days after the plating and continued until week 7. The electrode selection was based on the firing rate, as inferred from a prior Activity Scan (sparse 7x configuration). At each time point, a Network Assay (10 mins) and an Axon Scan (2 mins per configuration) were recorded to capture the network activity and allow for the inference of high-resolution electrical footprints (EFs) of individual neurons. At days in vitro (DIV) 52, the effect of 6-Cyano-7-nitroquinoxaline-2,3-dione (CNQX) on the firing rate of NGN2-iNs was assessed by recording 10 min of baseline activity and subsequently adding 20µM CNQX (Tocris) to each culture. Networks were allowed to equilibrate for 10 mins and another 20 min of network activity were recorded. Recordings were then concatenated to assess firing rate differences at the single-cell level. HD-MEA recordings were spike sorted using SpikeInterface (68) and Kilosort 2.5 (69) with default parameters and a spike threshold of 5.5. We used DeePhys (70) with default parameters to infer and aggregate electrophysiological features across the development (all feature groups), and calculated the UMAP embedding for each recorded culture. Single-cell UMAP embeddings were generated from all activity-based features at day 12, and clustering was performed using the Louvain algorithm. EFs were inferred from spike-sorted Axon Scans using custom-written Python code.

### Immunostaining

Patterned NGN2-iNs, co-cultured with rat cortical astrocytes (Genlantis) on PLO/LMN-coated 8-well or 96-well polymer coverslips (µ-Slide ibiTreat, ibidi) for 6 weeks, were fixed with 4% PFA for 10 mins at room temperature. Postfixation, the NGN2-iNs were quenched with 0.2 M Glycine (Sigma-Aldrich) in PBS for 20 mins and permeabilized with 0.05% Triton X100 (Sigma-Aldrich) in blocking buffer (BB, composed of PBS, 0.05% Tween-20 (Sigma-Aldrich), and 10% serum matching the secondary antibody’s species) for 10 mins at room temperature. Subsequently, the coverslips were incubated with BB (without Triton X100) for an additional 40 mins and then treated with the respective primary antibody diluted in BB for 2 hours at room temperature or at 4°C overnight. Following five washes with PBS for 5 mins each, the coverslips were incubated with the respective secondary antibody diluted in BB containing 1:1000 diluted DAPI for 1 hour at room temperature (protected from light). After another five washes with PBS for 5 mins each, the coverslips were stored in PBS at 4°C until image acquisition. The immunostained NGN2-iNs were imaged as confocal Z-stacks using the CSU-W1-SoRa inverted spinning disk confocal system, and the acquired images were processed with Fiji. Antibodies against the following proteins were purchased from the indicated vendors: MAP2 (EMD Millipore AB5622); SLC17A6 (Alomone Labs AGC-036-GP); SLC5A7 (Invitrogen PA5-117124); LHX9 (Sigma-Aldrich HPA009695); TRPM8 (Alomone Labs ACC-049), NTRK1 (FabGennix International TRKA-101Y).

## Data and code availability

The data generated for this study will be made available through NCBI’s Gene Expression Omnibus. All code generated in the study, including analysis parameters, will be made available at https://github.com/quadbio.

## Material availability

All experimental materials are available upon request to

## Supplementary Information

**Figure S1.**
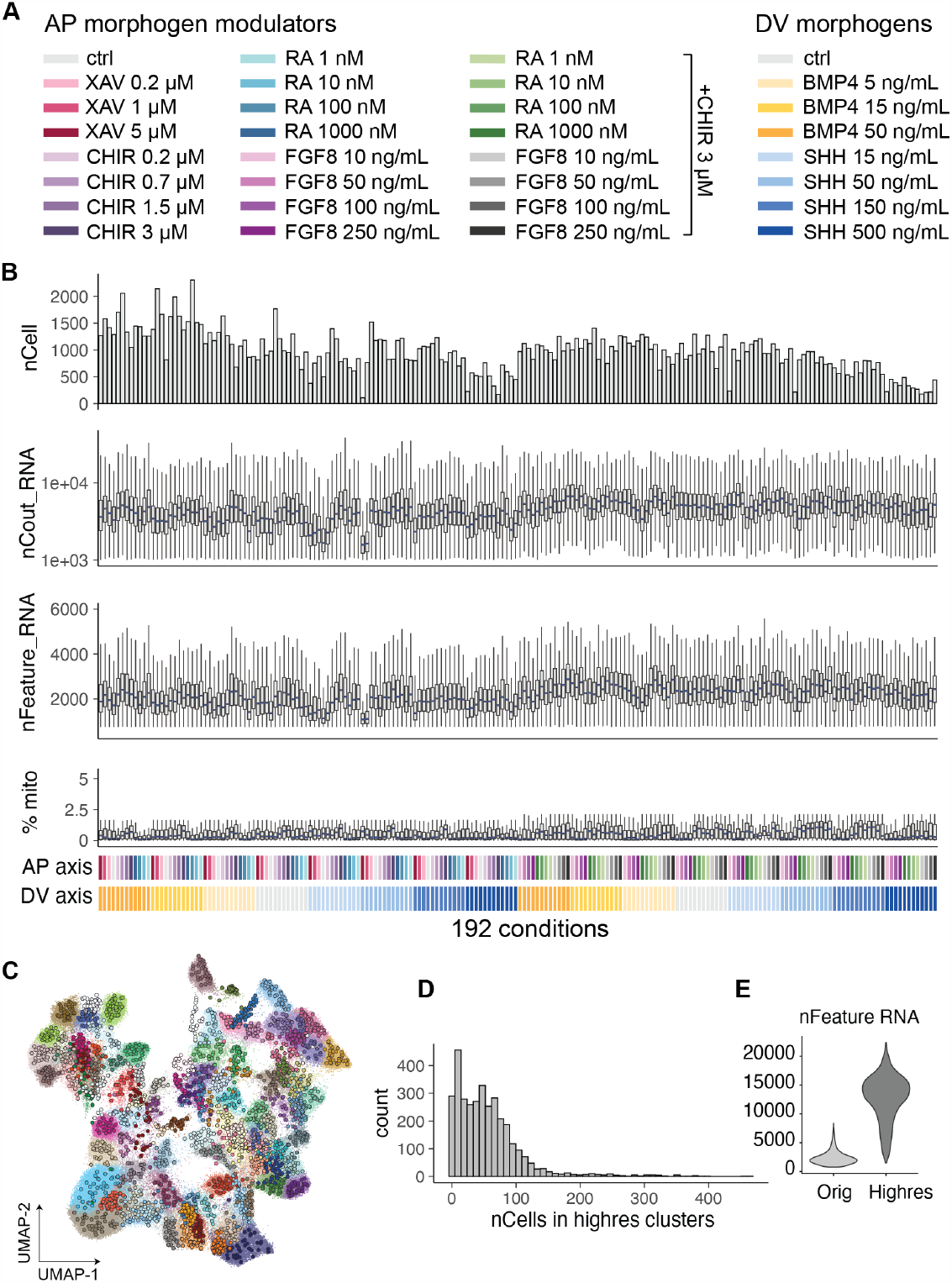
Experimental conditions and quality control of snRNA-seq for NGN2-iN combinatorial patterning screen. (A) Concentrations and color code for AP and DV morphogen signaling modulators used for combinatorial patterning. (B) Quality control of the snRNA-seq data in 192 morphogen combination conditions. Each column is a morphogen combination condition as color coded at the bottom. nCell, number of cells; nCount_RNA, total number of RNA molecules detected within a cell; nFeature_RNA, total number of genes detected within a cell; %mito, percentage of mitochondrial genes. (C) Overlay of UMAP embedding for the original single-cell dataset and high resolution clusters. (D) Distribution for the number of cells in each high resolution cluster. (E) Number of genes detected in the original data and high resolution clusters.

**Figure S2.**
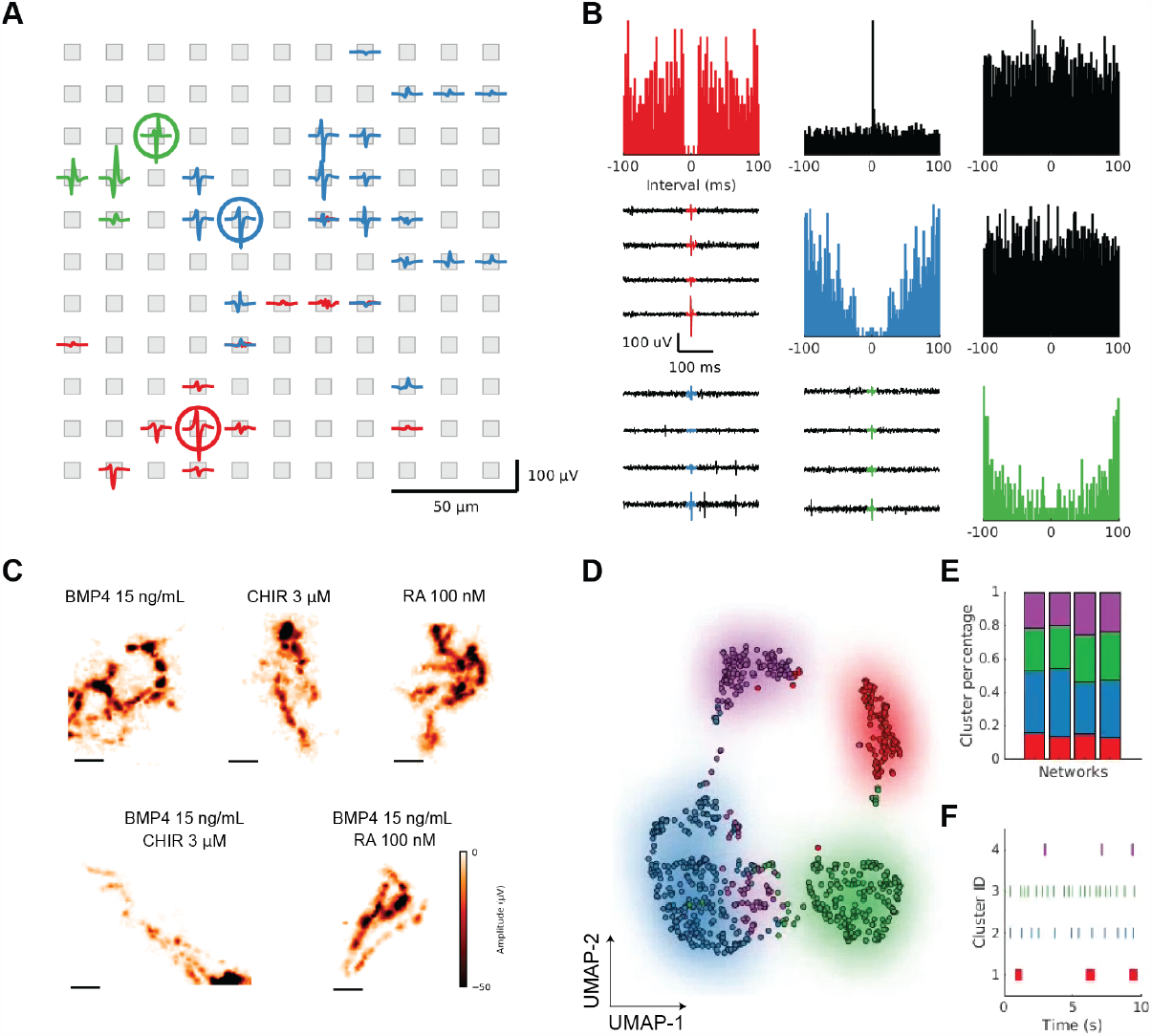
Electrophysiological recordings of patterned NGN2-iNs. Induced neurons developed spontaneous electrical activity two weeks after their plating on the high-density microelectrode arrays (HD-MEAs), and were recorded for up to three months. (A) Spike-triggered electrical footprints (EFs) of three example spike sorted units (i.e., putative neurons). Red, blue and green colors indicate the different unit IDs. (B) Example spike-train autocorrelograms (diagonal), and cross-correlograms (upper triangle, in black), for the three example units from panel (A). (C) EFs indicated robust outgrowth of neurites on the HD-MEA in all patterning conditions at day 12. Colors indicate the average amplitude of spike-triggered EFs recorded on the HD-MEA. Scale bar: 200 µm. (D) Example UMAP embedding of 798 units pooled over n=4 networks (CHIR+BMP4 patterning), using the activity-based features from their day 12 recording as input. The coloring indicates the cluster ID obtained from a Louvain clustering. (E) The overall distribution of the obtained clusters was comparable across the four different networks; see (D) for information on colors. (F) Example spike trains for each of the inferred clusters indicate clear differences in the activity of each obtained cluster. Each line represents one action potential; see (D) for information on cluster colors.

**Figure S3.**
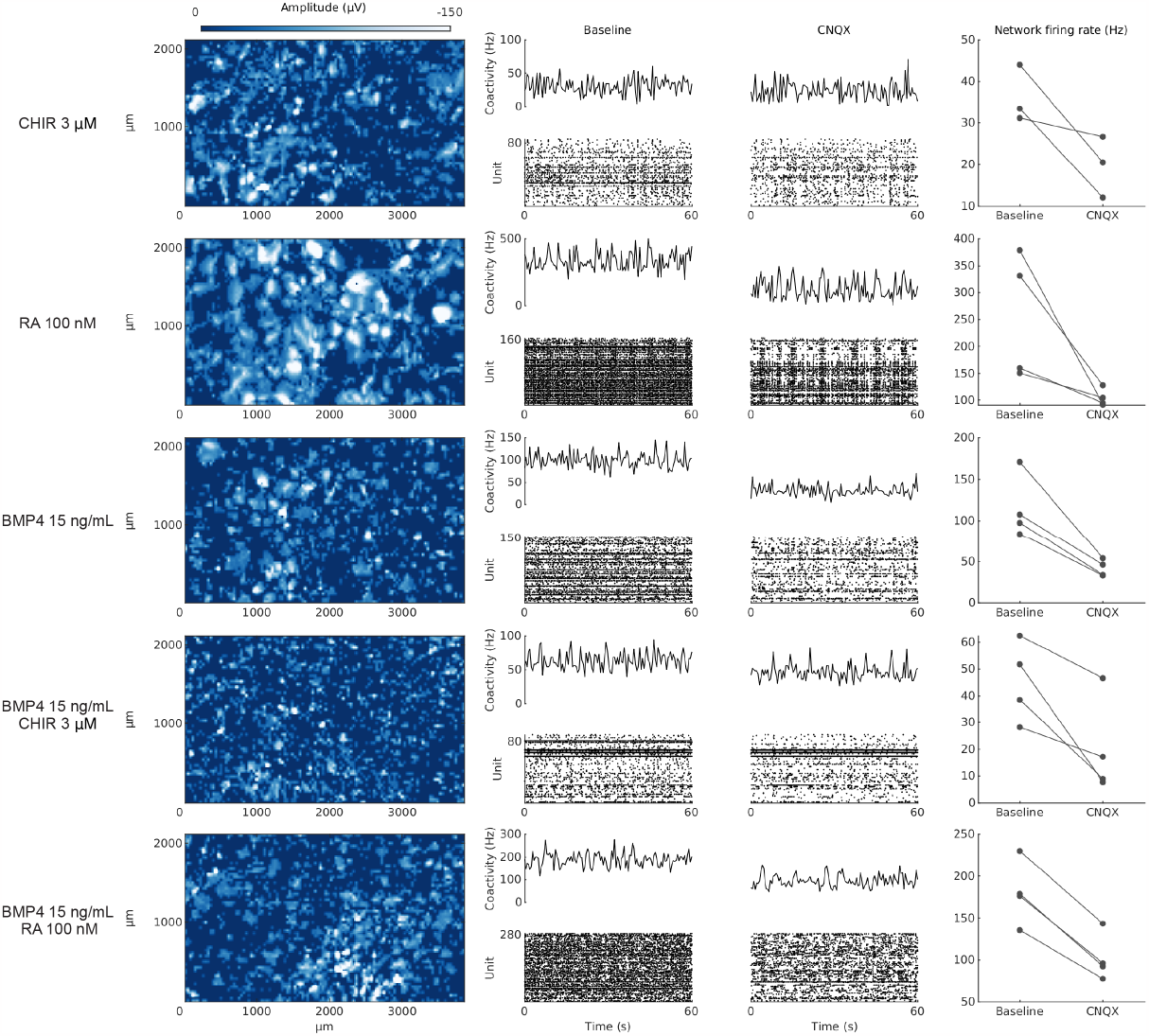
Spontaneous activity of patterned NGN2-iNs is altered by CNQX. Column on the left: Example whole-array spike amplitude maps for five patterning conditions indicating the distribution of neurons on five different HD-MEAs; colors indicate the average spike amplitudes as obtained during the Activity Scan for each HD-MEA. Middle column: Example spike raster plots indicating the ongoing network activity during a baseline recording and following the application of CNQX (only 60 seconds are shown for each condition; the CNQX recording was recorded 10 mins after the drug application). Column on the right: The application of the AMPA receptor antagonist CNQX significantly reduced the mean network firing rate; each dot indicates one measured network (experiments were performed at day 45).

**Figure S4.**
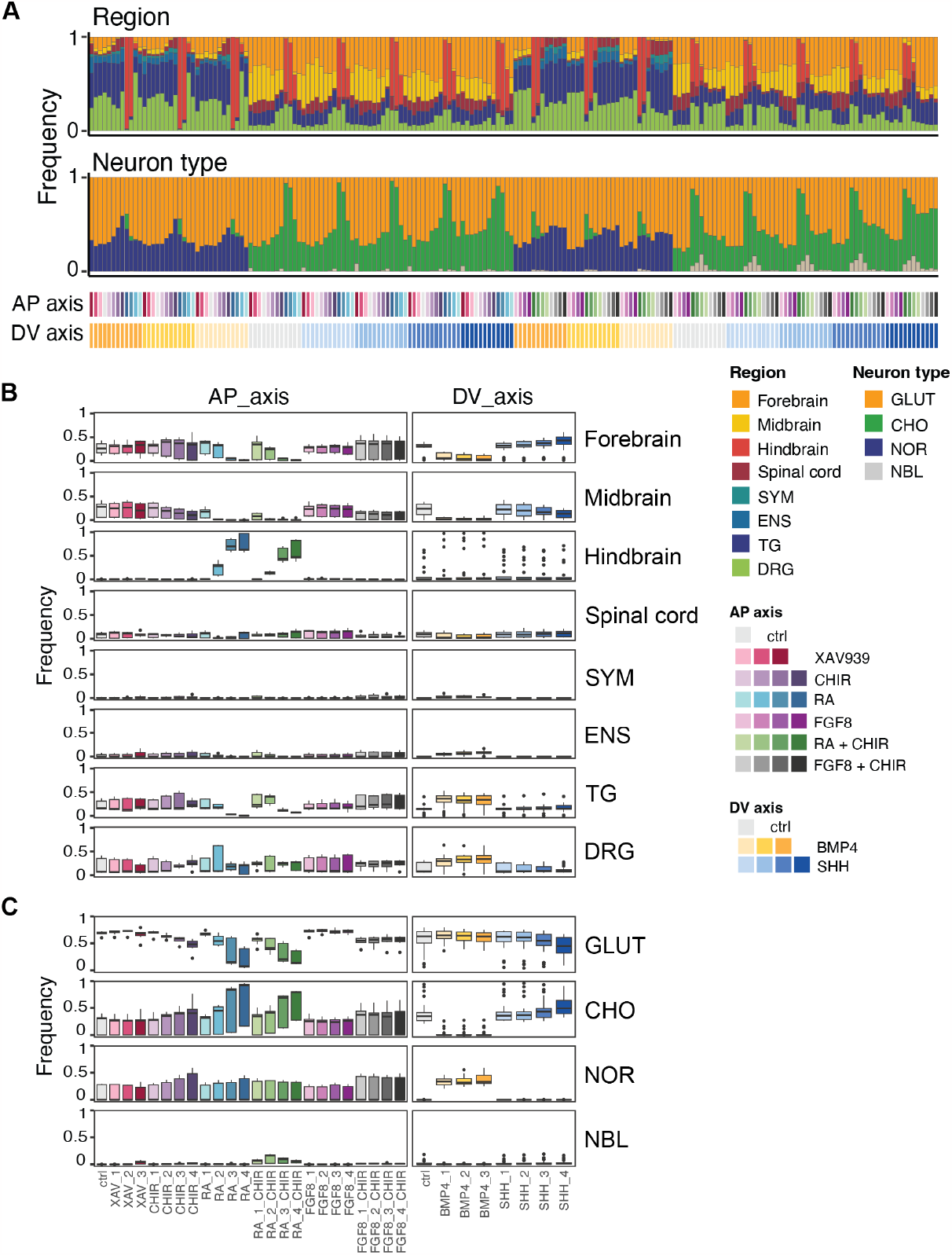
Contribution of single and combinatorial morphogen signaling modulators to patterned NGN2-iN diversity. (A) The stacked bar plot illustrates the distribution of cells among annotated regional identities and neuron types within each morphogen combination. Each column represents a specific morphogen combination condition, color-coded as depicted at the bottom. Also refer to Fig. S1A for the respective morphogen signaling modulator concentrations associated with each color. SYM, sympathetic nervous system; ENS, enteric nervous system; TG, trigeminal ganglia; DRG, dorsal root ganglia; GLUT: glutamatergic neuron; CHO, cholinergic neuron; NOR, noradrenergic neuron; NBL, neuroblast-like cells. (B-C) Assessing the impact of morphogen signaling modulators on regional identity (B) and neuron type (C) composition. Each column represents a distinct concentration of AP or DV morphogen signaling modulator. The boxplot shows the frequency of annotations (indicated in each row) in samples that has this specific concentration of AP or DV morphogen signaling modulator.

**Figure S5.**
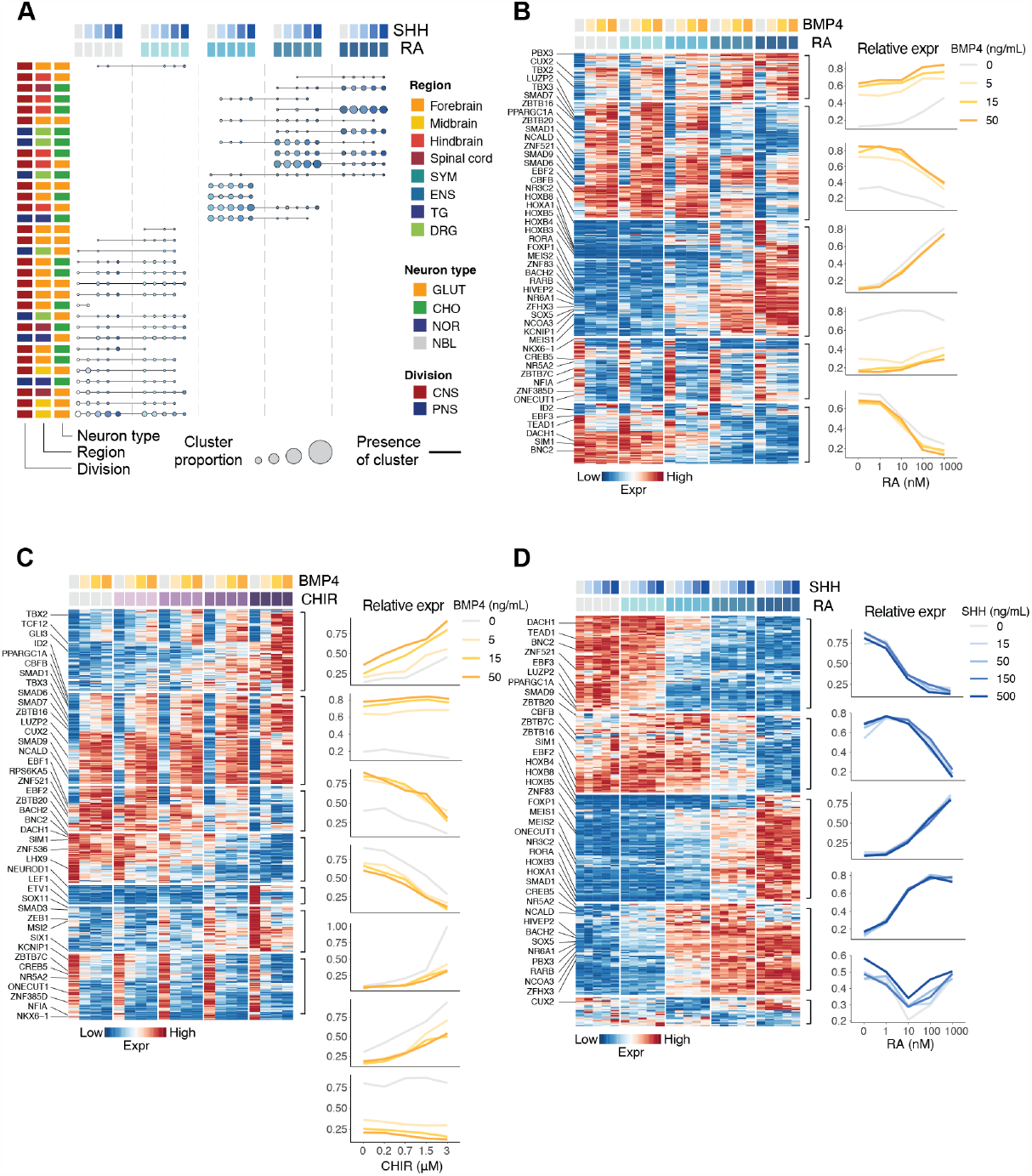
Unfolding the gene programs coupling neural identity shift in response to morphogen combinations. (A) Dot plot depicting cluster composition and identity shift in response to morphogen combination gradients. Similar to Fig. 2B. (B-D) Molecular changes underlying morphogen combination gradients. Heatmap (left) rows (only transcription factors are labeled) represent genes identified by random forest classifier for its association with morphogen-induced gene expression changes. These genes are further grouped into gene programs based on similarity of expression patterns. Columns colored by condition. Line plots (right) show average expression of genes within each gene program visualized across morphogen combination gradients.

**Figure S6.**
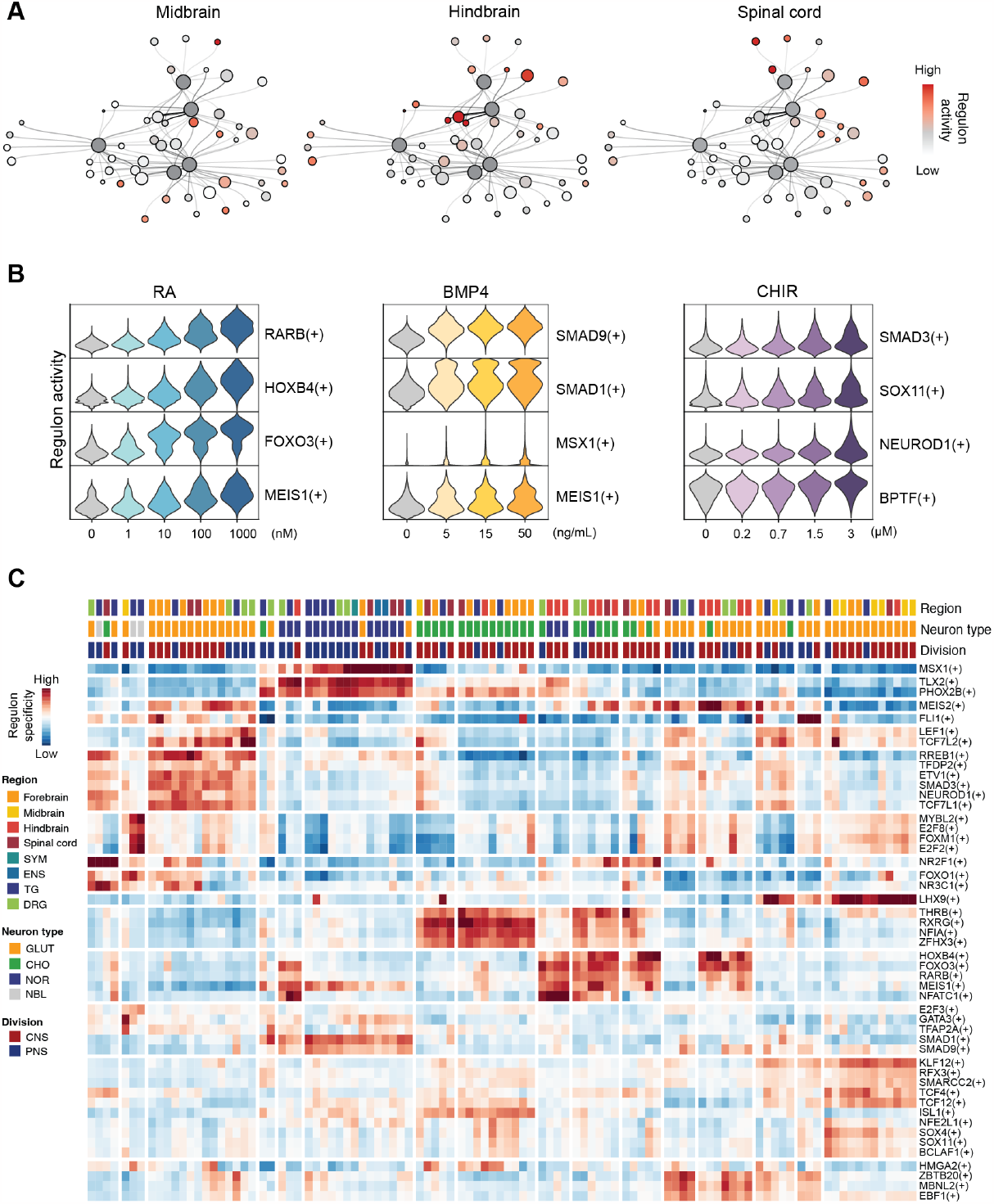
Analysis of regulons inferred from combinatorial patterned NGN2-iNs. (A) Gene regulatory network (GRN) centered on morphogen signaling modulators (depicted in dark grey) with nodes colored based on the activities of corresponding regulons within subsetted cells in each indicated regional identity (midbrain, hindbrain and spinal cord). (B) Violin plots illustrating the activities of indicated regulons under each concentration of the morphogen signaling modulator. The color code corresponds to the identities and concentrations morphogen signaling modulators as illustrated in Fig. S1A.

**Figure S7.**
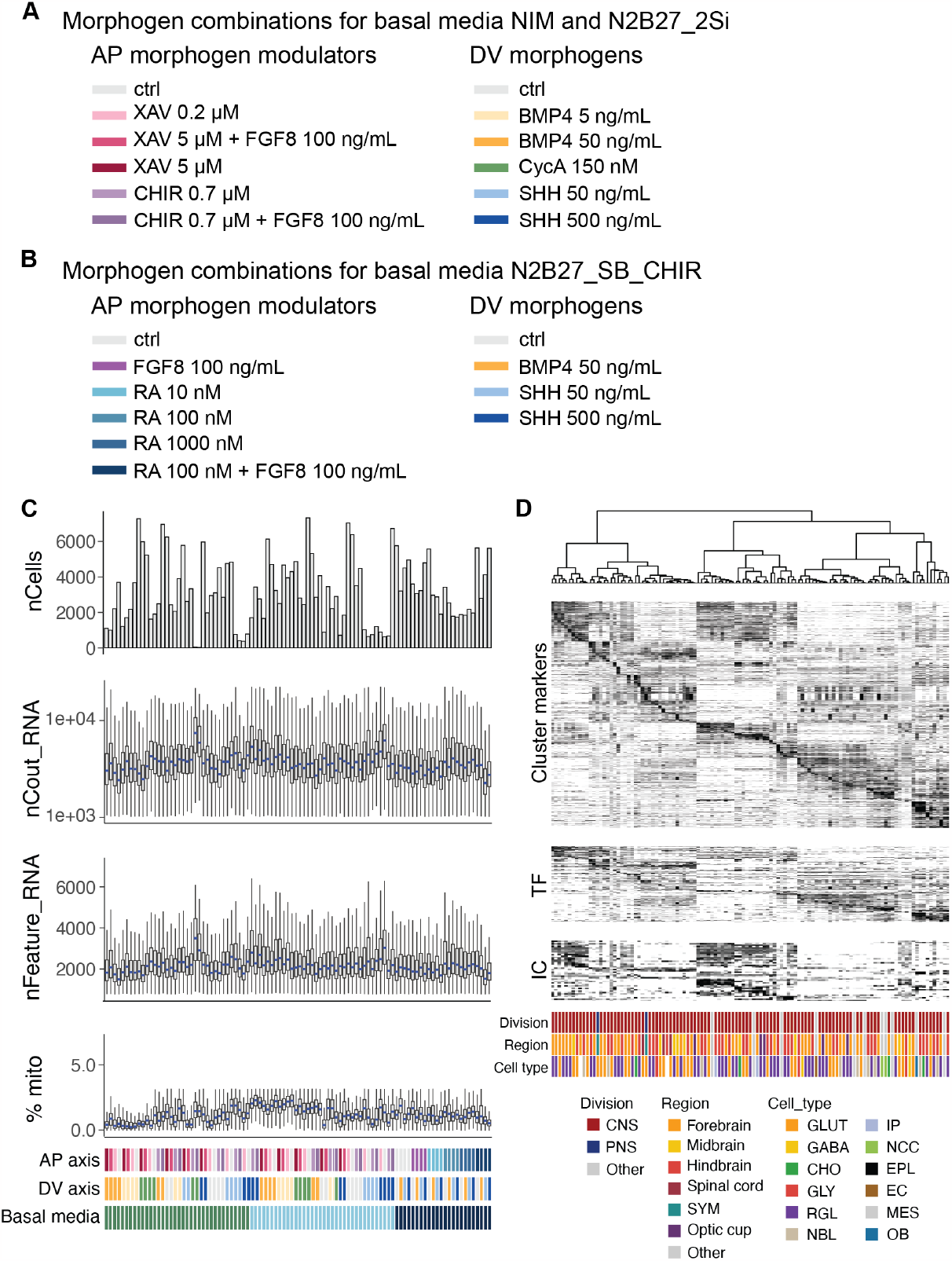
Experimental conditions and quality control of snRNA-seq for pre-patterned NGN2-iNs. (A-B) Concentrations and color code for AP- and DV- morphogen signaling modulators coupled with basal media NIM, N2B27-2Si (A) and N2B27-SB-CHIR (B). (C) Quality control assessment of snRNA-seq data obtained from the pre-patterning screen. Each column represents a unique condition, color-coded as depicted at the bottom. Refer to (A) for the respective concentrations of each morphogen signaling modulator. (D) Heatmap showing cluster marker genes, transcription factors (TFs) and ion channels (ICs) expression in each pre-patterned NGN2-iN cluster. The annotation of each cluster, including division, region, and cell type, is depicted with color-coded sidebars. GABA, gabaergic neuron; GLY, glycinergic neuron; RGL, radial glia-like cell; IP, intermediate progenitor; NCC, neural crest cells; EPL, ependymal-like cell; EC, endothelial cell; MES, mesenchymal cell; OB, osteoblast.

**Figure S8.**
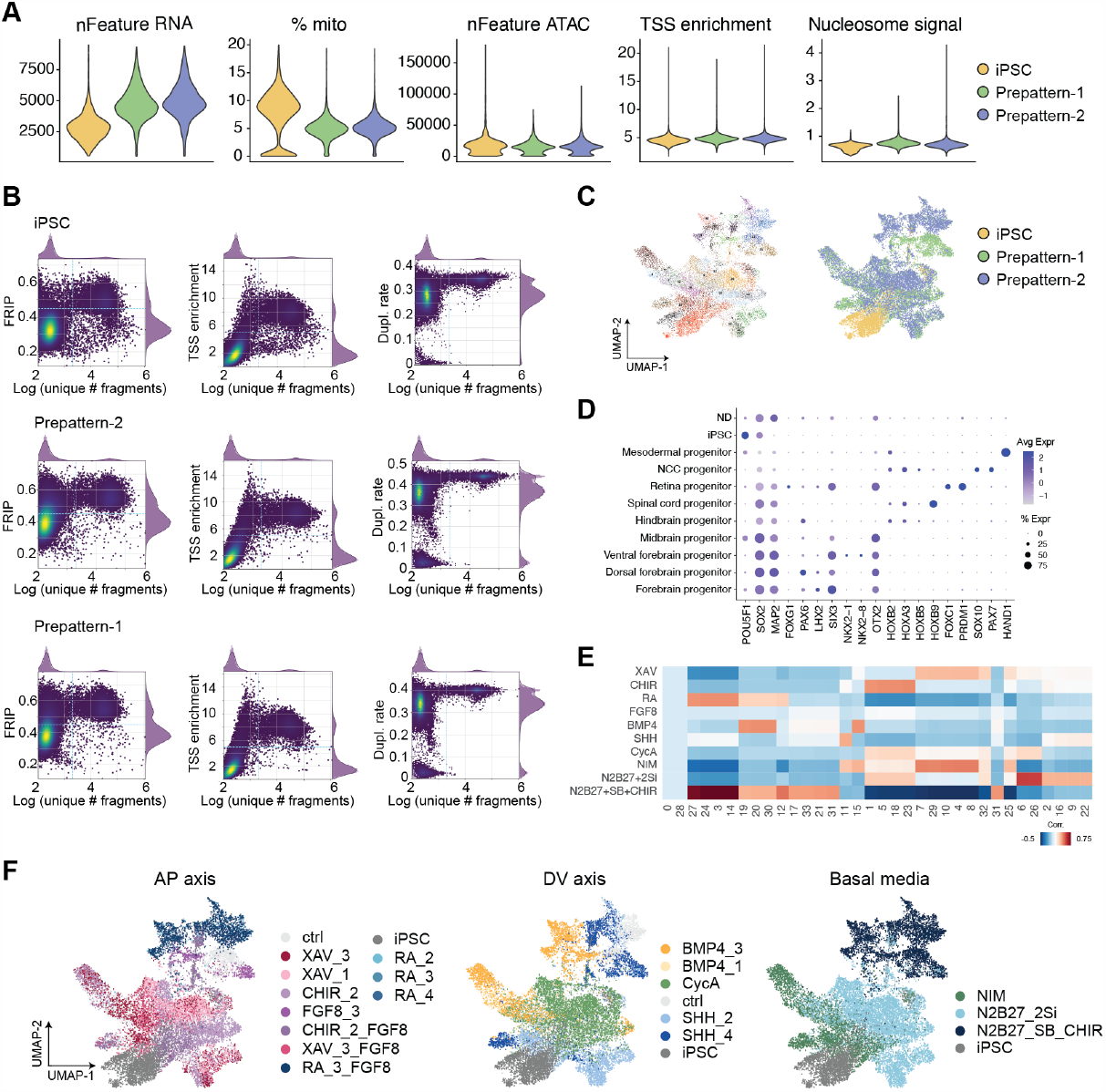
Quality control and supplemental multiome analysis of pre-patterned neural stem cells and progenitors. (A) Quality assessment of the multiome data. Three 10x multiome experiments are run independently and integrated, including iPSC and two other reactions for pre-patterned progenitor cells. Sample composition in each reaction is shown in Supplementary Table 1. TSS: transcription start site. (B) Scatter plots of quality metrics and used thresholds during filtering. FRIP: fraction reads in peak. (C) UMAP embedding of the integrated data, showing final clustering based on transcriptomes as well as the original identity of samples. (D) Dotplot showing marker gene expression used to annotate samples. (E) Demultiplexing of multiome data from bulk RNA-seq using MUSiC. Heatmap shows correlation of predicted bulk composition with bulk conditions. 0 and 28 are clusters from control sample and were not predicted in any bulk sample. (F) UMAP embeddings are colored by the demultiplexed AP, DV and basal media identities.

**Figure S9.**
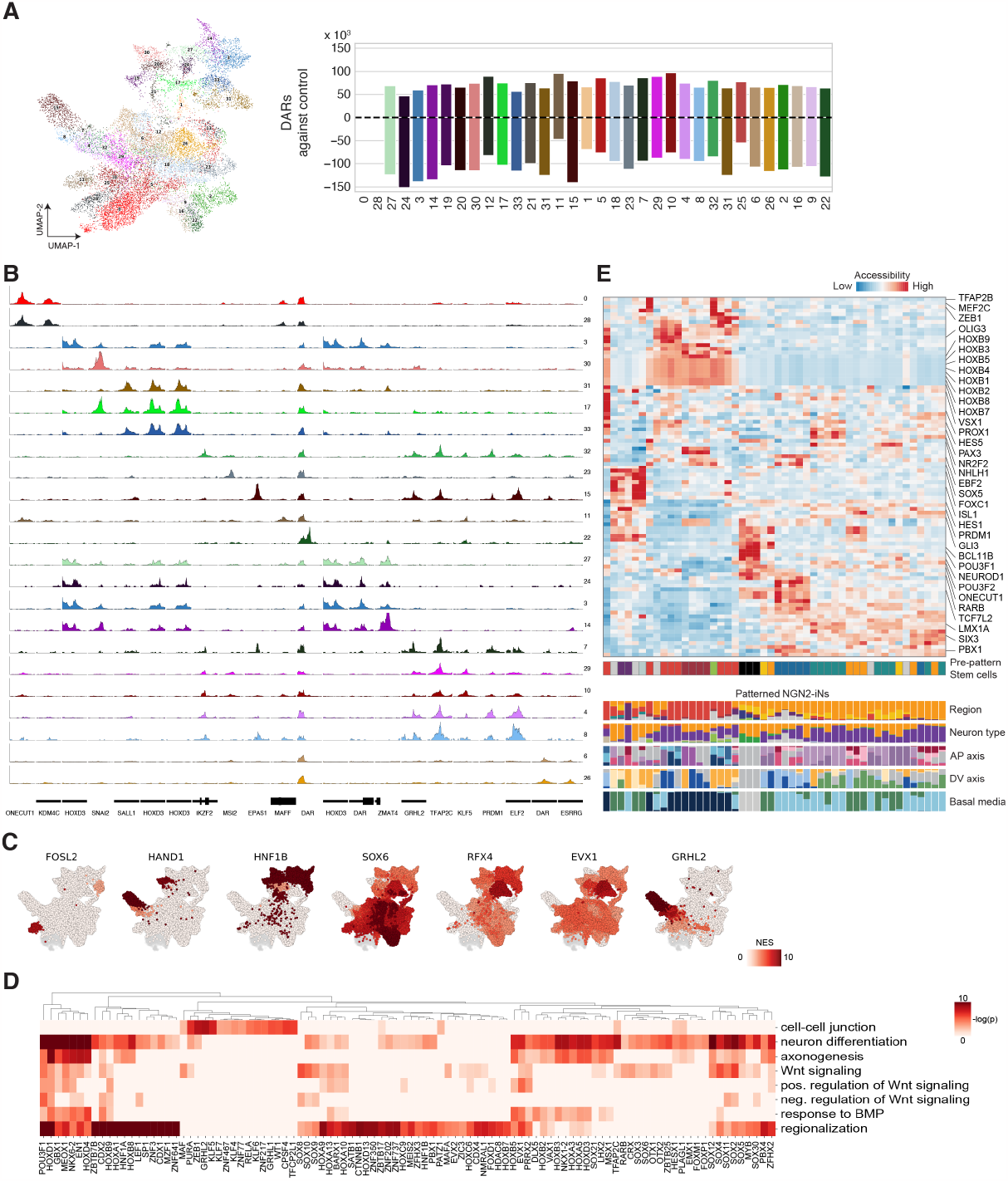
Quality control and supplemental multiome analysis of pre-patterned neural stem cells and progenitors. (A) UMAP embedding of the integrated data, showing final clustering (left). Barplots showing the number of differentially accessible regions DARs (pval<1e-10, lfc>2) of each cluster against the two control clusters (0, 28), showing large-scale closing and opening of regions upon morphogen exposure (right). (B) Chromating landscape of clusters for different neuronal regions linked to TF expression. (C) UMAP embedding colored by motif enrichment in accessible regions of that cluster (normalized enrichment score, NES). (D) Heatmap showing GO enrichment of genes linked to regions part of the cistrome of different TFs. (E) Heatmap showing differential accessibility of NGN2-motifs and associated TFs in pre-patterned neural stem cell and progenitor clusters (top) and compositions of the resulting pre-patterned NGN2-iNs from corresponding neural stem cell clusters in (bottom).

**Figure S10.**
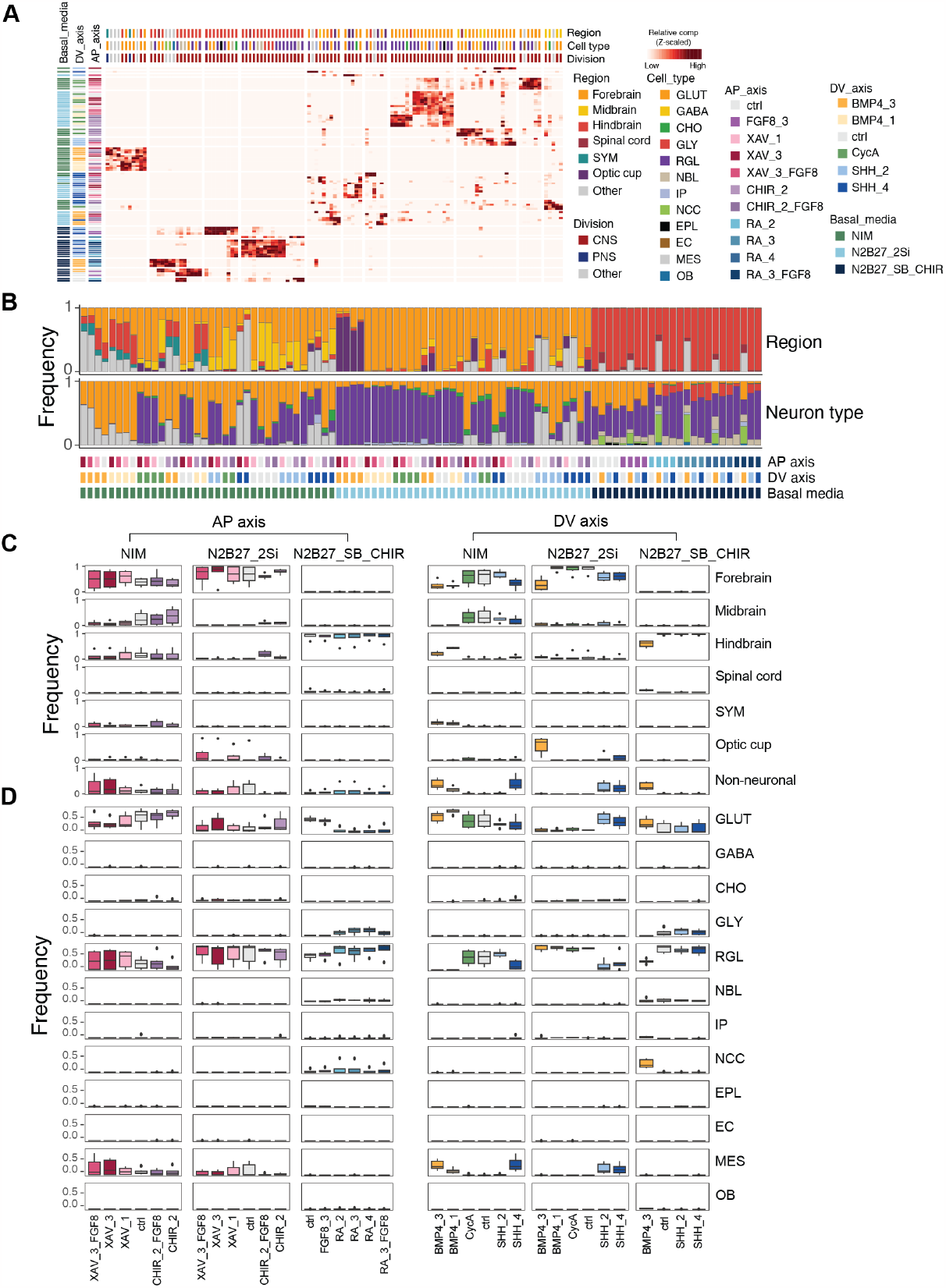
Interplay between morphogen combinations and basal media contributes to pre-patterned NGN2-iN diversity. (A) Heatmap visualization of the contribution of each combinatorial morphogen condition (row) to pre-patterned NGN2-iN clusters (column). The cell count within each morphogen condition is normalized for comparative analysis. Color-labeled sidebars denote morphogen conditions and annotations of cell clusters. (B) Composition of regional identity and cell type in distinct AP-DV-basal media morphogen signaling modulator conditions. Similar to Fig. S4A. (C-D) The effect of basal media and morphogens on neural identity (C) and cell type (D) composition in each of the indicated AP or DV morphogen signaling modulator concentration. Similar to Fig. S4B to C.

**Figure S11.**
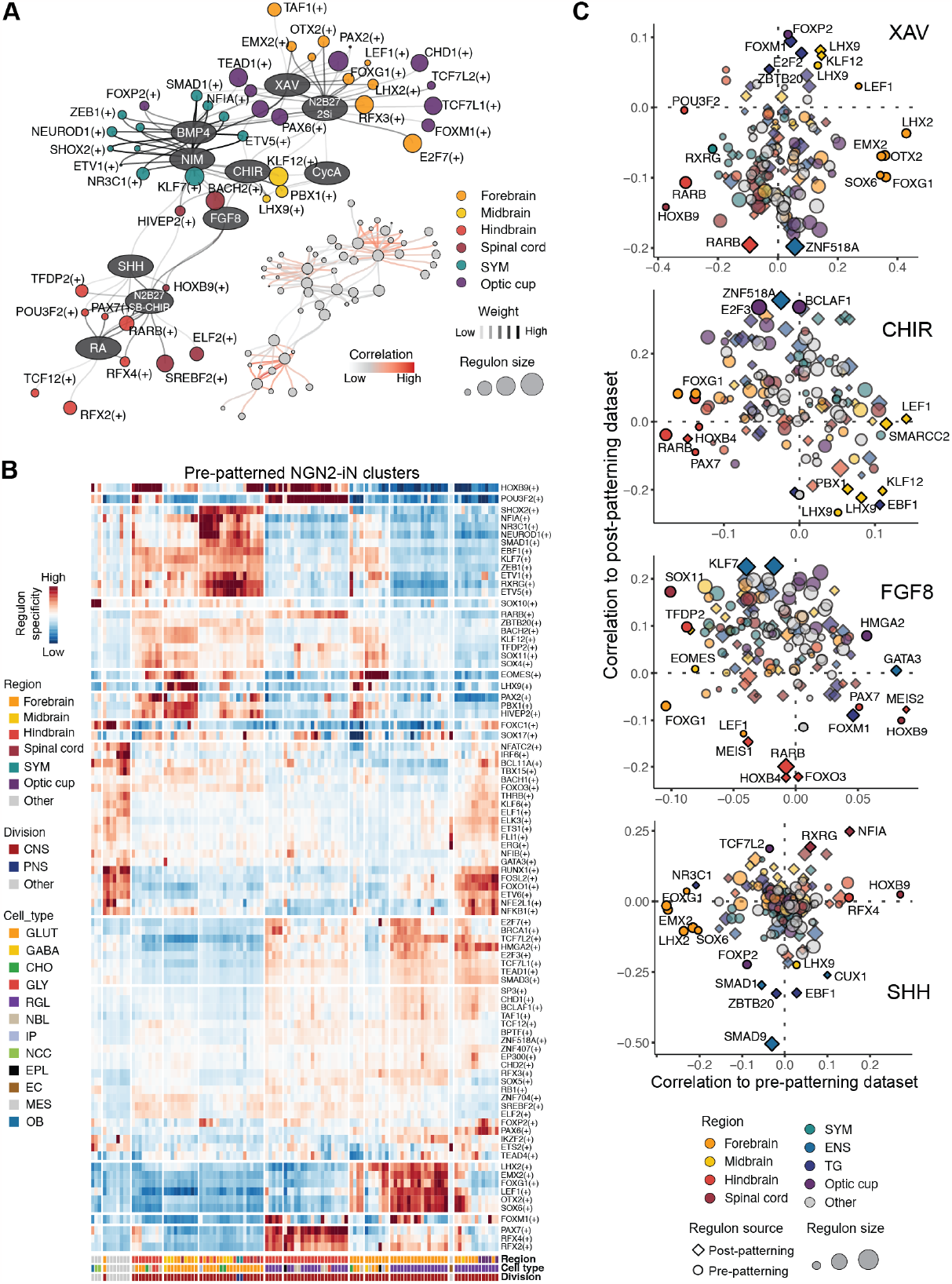
Gene regulatory network and regulon analysis of pre-patterned NGN2-iNs. (A) Morphogen signaling modulators and basal media centered gene regulatory network with regulons inferred from pre-patterned NGN2-iNs. Similar to Fig. 2F. On the left side, each oval node denotes a morphogen signaling modulator linked to circular nodes indicating associated regulons. Edge colors signify the strength of the association between morphogen signaling modulators and regulons. The regulons are color-coded based on the regional identity in which they exhibit maximum activity. Node sizes correspond to the regulon sizes, reflecting the number of target genes regulated by the TF. On the right side, the same GRN is depicted, where edge color represents the correlation of regulon activity with morphogen modulator concentration (B) Heatmap depicting regulon specificity in pre-patterned NGN2-iN clusters. The annotation of each pre-patterned NGN2-iN cluster is color coded as side bars. (C) Comparison of regulons in pre- and post-patterned NGN2-iNs. Similar to Fig. 3L.

**Figure S12.**
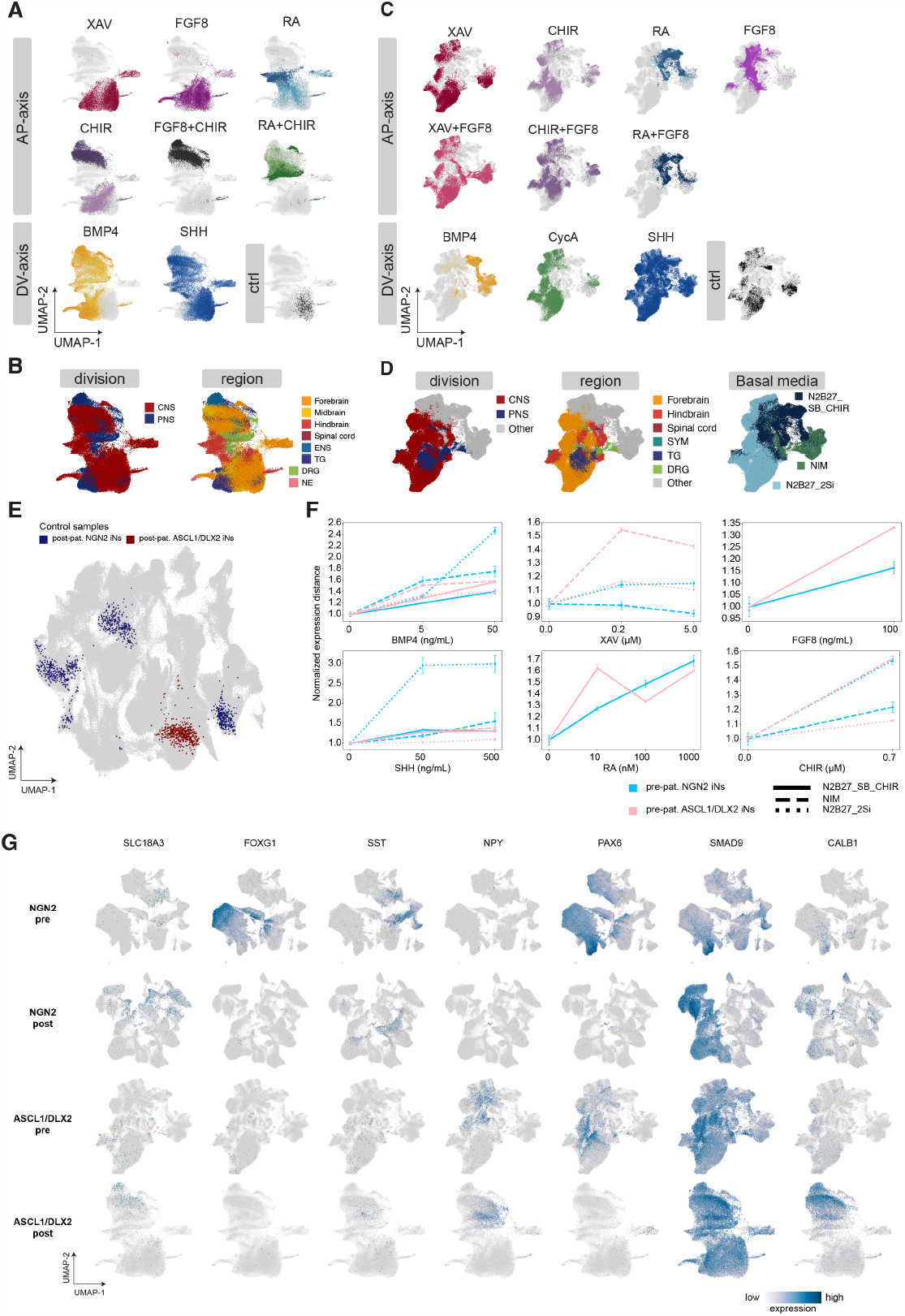
Influence of morphogen combinations and basal media on post- and pre-patterned ASCL1/DLX2-iNs diversity. (A-B) Annotation of post-patterned ASCL1/DLX2-iNs to different brain regions. UMAP embedding of 85,756 cells in the prepatterned ASCL1/DLX2-iN dataset, colored by cluster identity, the source of AP- or DV-morphogens (A), the annotations transferred from primary neuron reference atlases and the basal media used (B). (C-D) Annotation of pre-patterned ASCL1/DLX2-iNs to different brain regions. UMAP embedding of 139,711 cells in the pre-patterned ASCL1/DLX2-iN dataset, colored by cluster identity, the source of AP- or DV-morphogens (C), the annotations transferred from primary neuron reference atlases and the basal media used (D). CNS, central nervous system; PNS, peripheral nervous system; ENS, enteric nervous system, TG; trigeminal ganglion; DRG, dorsal root ganglion; NE, neuroepithelium. (E) UMAP embedding of all neuronal cells across NGN2 and ASCL1/DLX2-iNs (n=430,936 cells). Control cells (mTESR medium, no morphogens) are colored. (F) Scatter plot showing mean distance of post-patterning NGN2-iNs and ASCL1/DLX2-iNs to control cells as a function of morphogen concentration. Error bars represent SEM. Different basal media are shown in different line styles. (G) UMAP embeddings showing gene expression in NGN2-iNs and ASCL1/DLX2-iNs.

**Figure S13.**
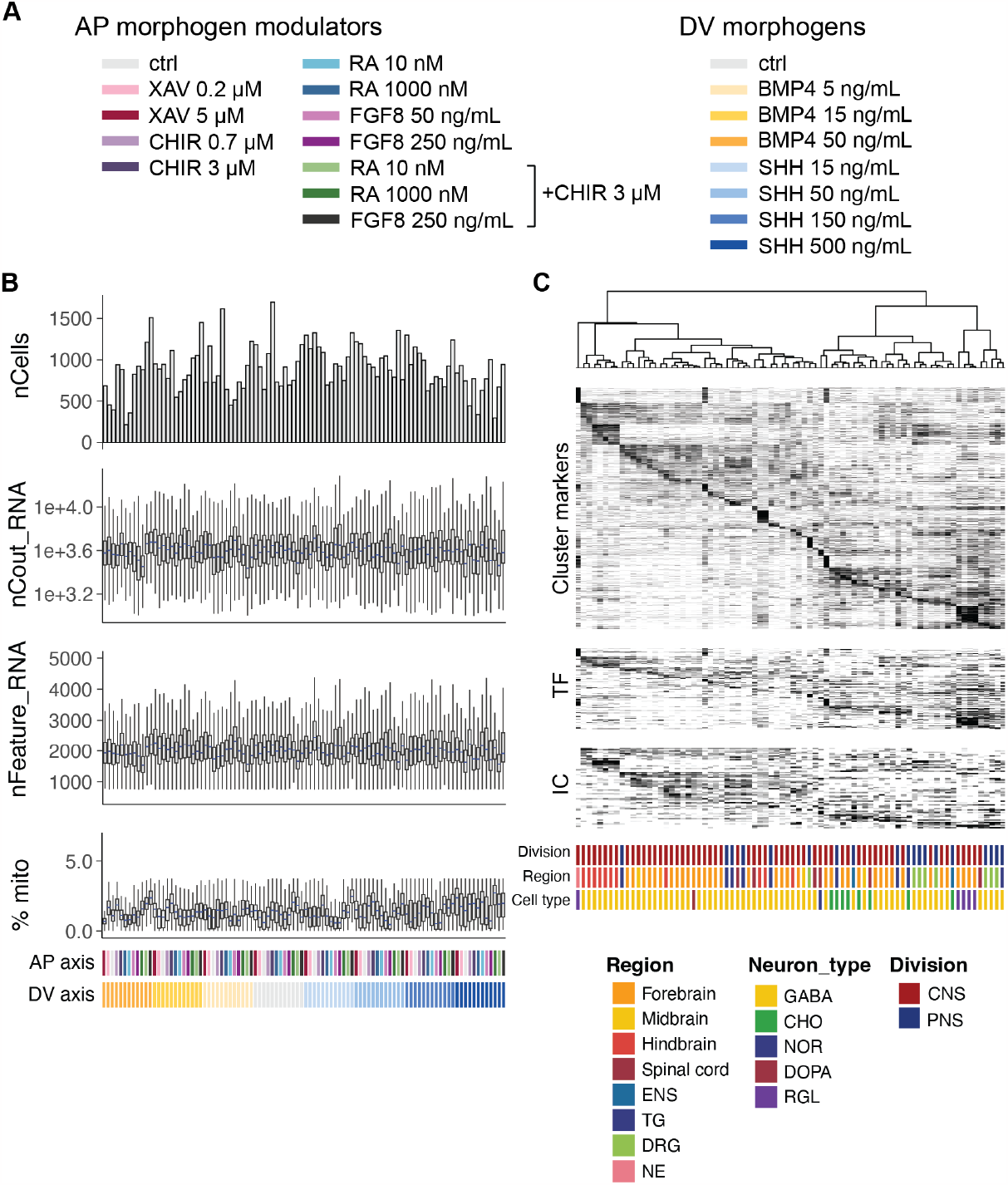
Experimental conditions and quality control of snRNA-seq for ASCL1/DLX2-iNs combinatorial post patterning screen. (A) Concentrations and color codes representing AP- and DV-morphogen signaling modulators utilized for combinatorial postpatterning of ASCL1/DLX2-iNs. (B) Evaluation of snRNA-seq data quality. Each column represents a morphogen combination condition color-coded at the bottom. (C) Heatmap illustrating the expression of cluster marker genes, transcription factors (TFs), and ion channels (ICs) in each post-patterned ASCL1/DLX2-iN cluster.

**Figure S14.**
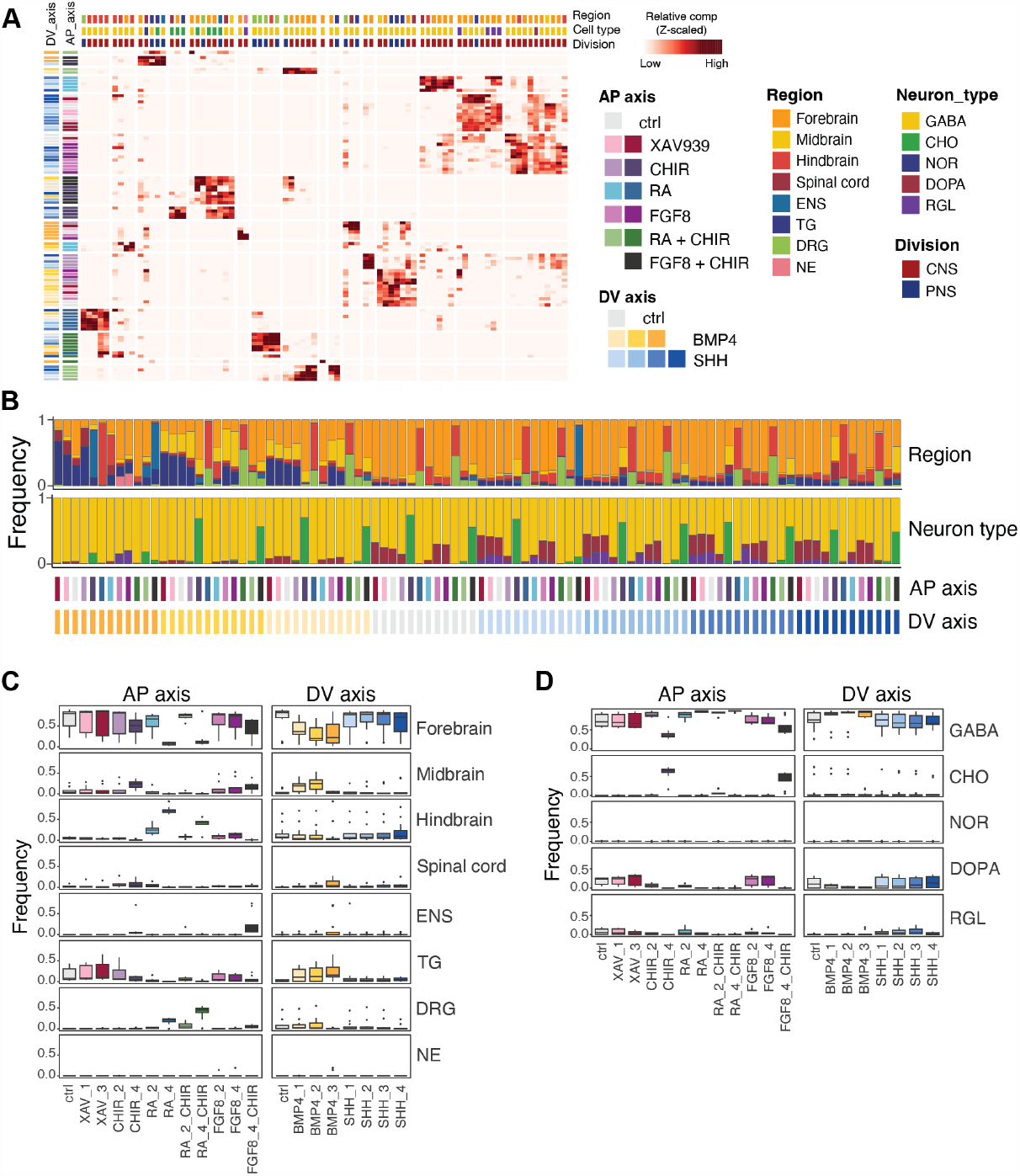
Contribution of single and combinatorial morphogen signaling modulators to post-patterned ASCL1/DLX2-iN diversity. (A) Depiction of cluster distribution (columns) in each post-patterned ASCL1/DLX2-iN sample (rows). AP and DV morphogen signaling modulator concentrations and cluster annotations are depicted as color-coded side bars. NE, neuroepithelium; GABA, gabaergic neuron; DOPA, dopaminergic neuron. (B) Overview of regional identity and cell type composition in each post-patterned ASCL1/DLX2-iN sample. (C-D) Evaluation of the impact of morphogen signaling modulators on regional identity (C) and neuron type (D) composition on post-patterned ASCL1/DLX2-iN neurons.

**Figure S15.**
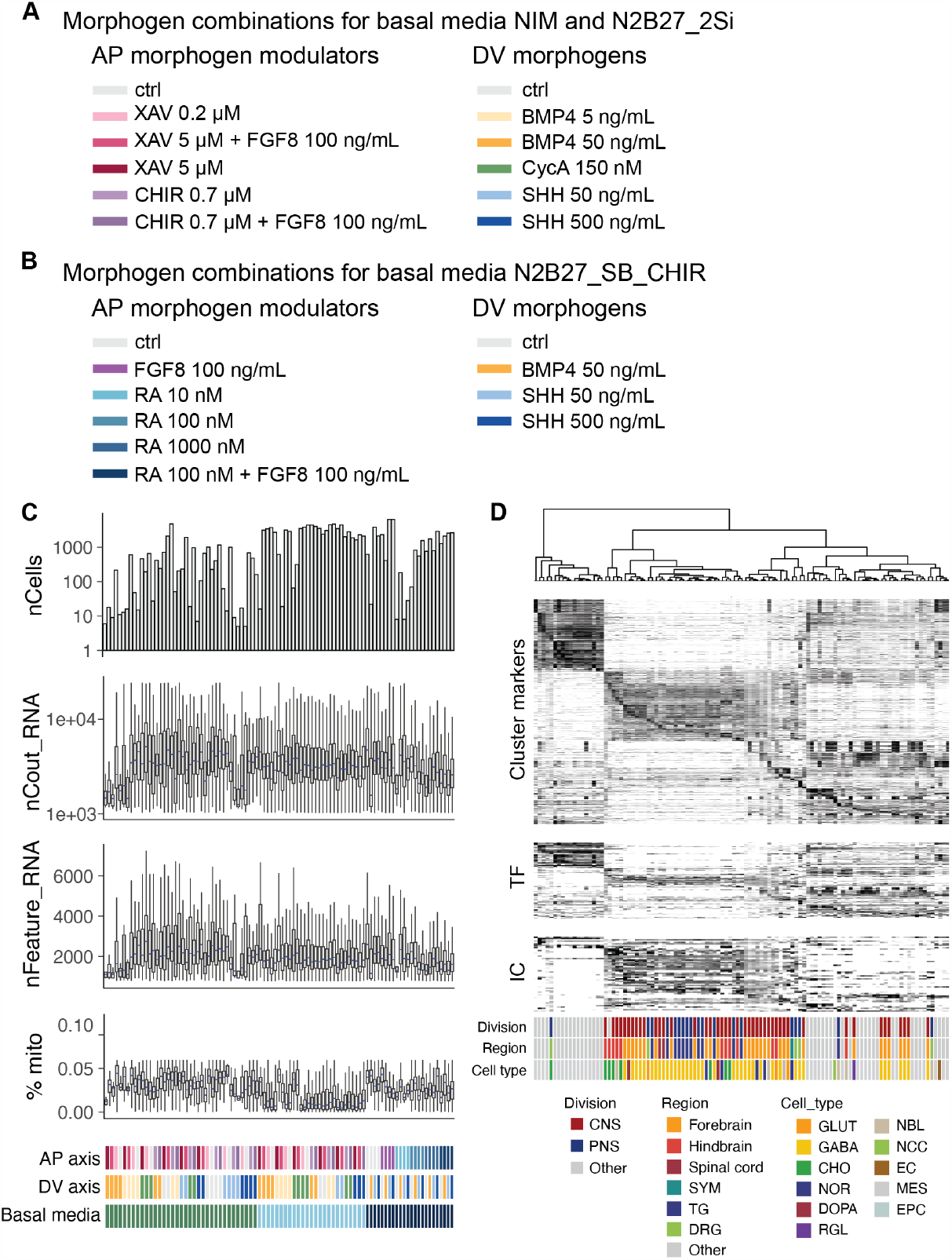
Experimental conditions and quality control of snRNA-seq for pre-patterned ASCL1/DLX2-iNs. (A-B) Depiction of concentrations and color coding for AP and DV morphogen signaling modulators coupled with basal media NIM, N2B27-2Si (A), and N2B27-SB-CHIR (B). (C) Evaluation of snRNA-seq data quality from the ASCL1/DLX2-iN pre-patterning screen. Each distinct condition is color-coded at the bottom. Refer to (A) for corresponding concentration of morphogen signaling modulators. (D) Heatmap illustrating the expression of cluster marker genes, transcription factors (TFs), and ion channels (ICs) in each pre-patterned ASCL1/DLX2-iN cluster.

**Figure S16.**
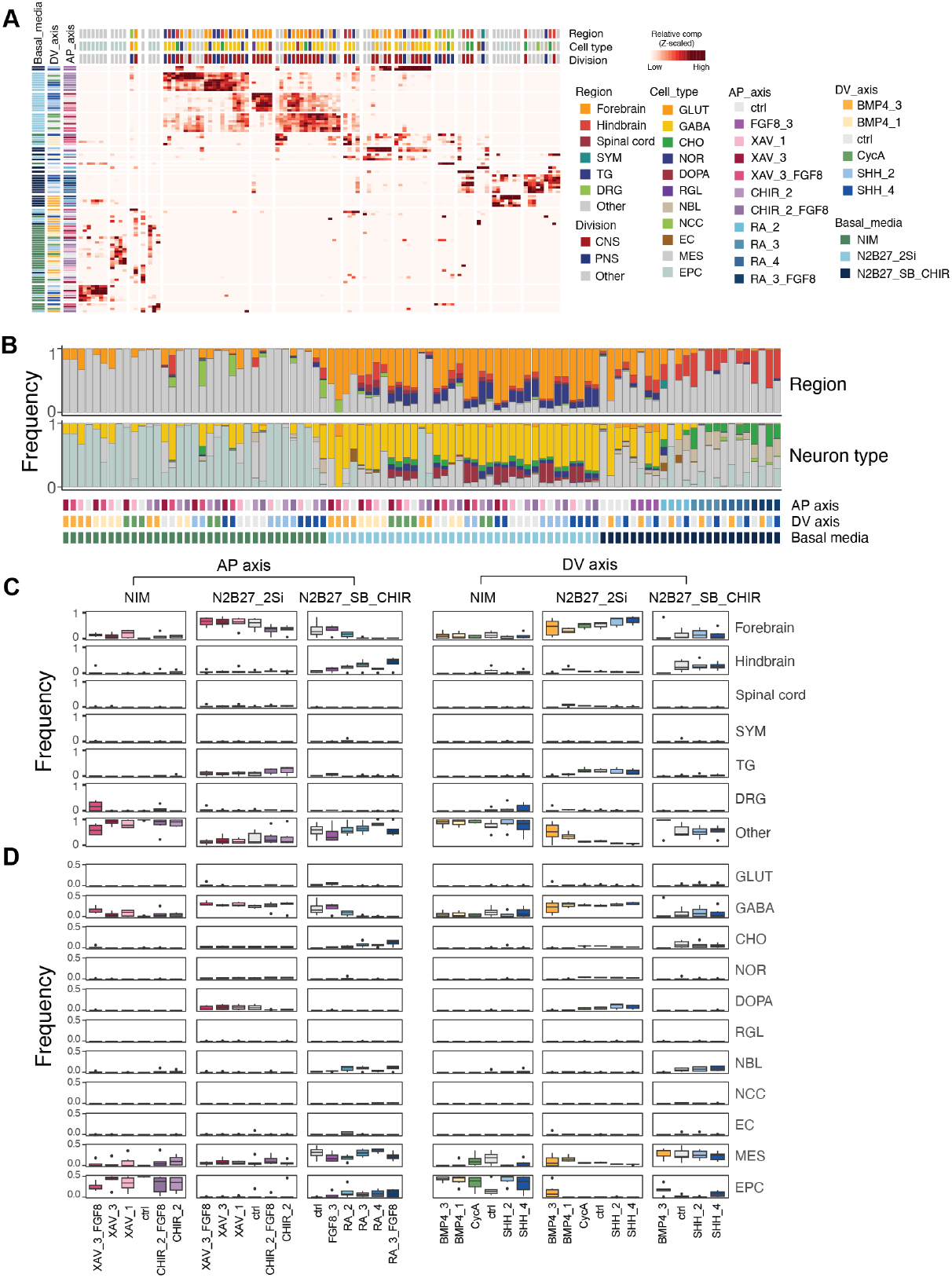
Influence of morphogen combinations and basal media on pre-patterned ASCL1/DLX2-iN diversity. (A) Representation of cluster distribution (column) in each pre-patterned ASCL1/DLX2-iN sample (row). EC: endothelial cell; MES: mesenchymal cell; EPC: epithelial cell. (B) Composition of regional identity and cell type in each pre-patterned ASCL1/DLX2-iN sample. (C-D) Impact of basal media and morphogens on regional identity (C) and cell type (D) composition.

**Figure S17.**
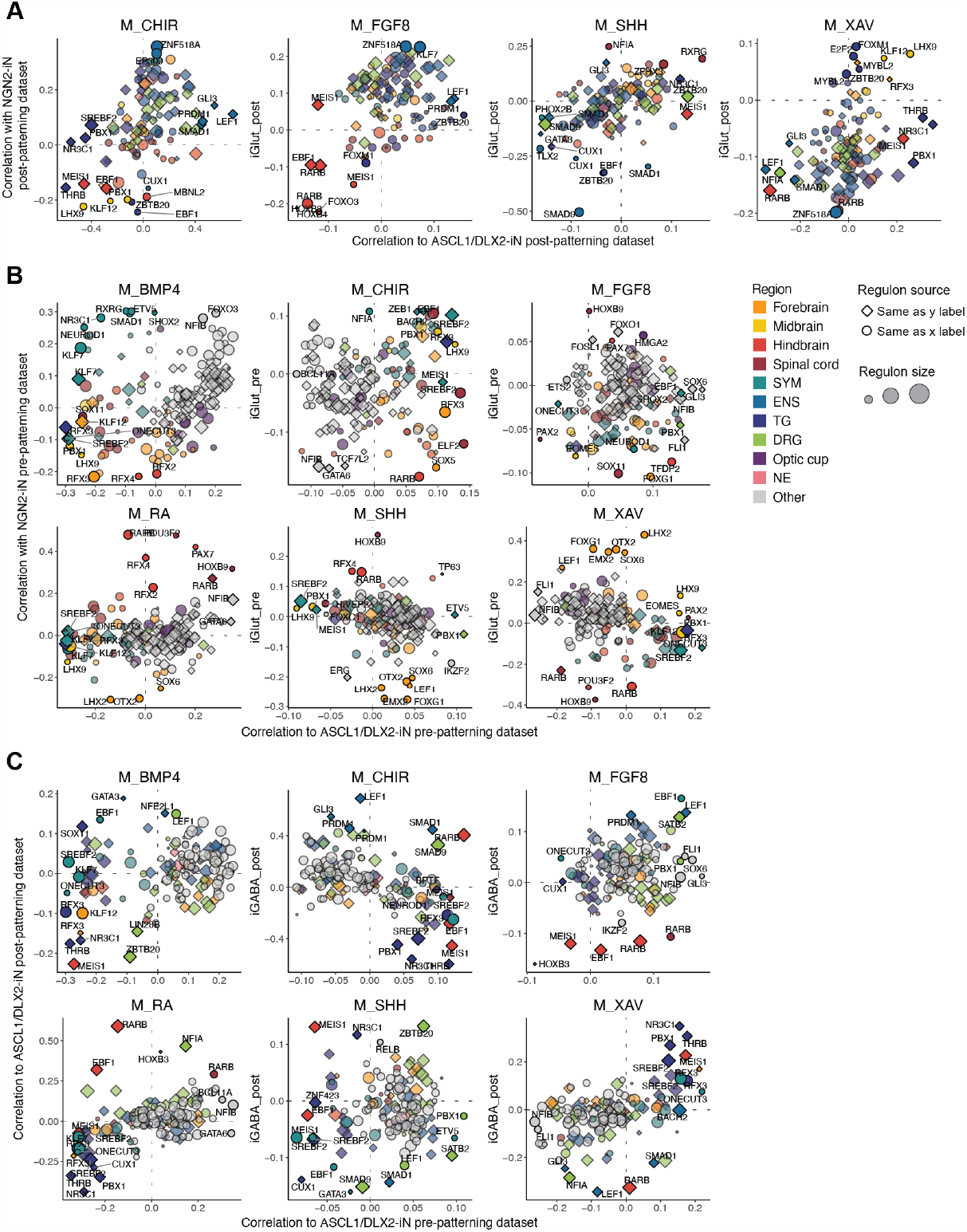
Regulon comparison of NGN2-iNs and ASCL1/DLX2-iNs. (A-C) Regulon comparison between (A) post-patterned NGN2-iNs and ASCL1/DLX2-iNs (B) pre-patterned NGN2-iNs and ASCL1/DLX2-iNs (C) pre- and post-patterned ASCL1/DLX2-iNs. Regulons are colored with the regional identity where maximal activity is registered, while the size of the regulons was represented as the size of the dots. Similar to Fig. 4F.

